# Closed-Loop Control of Anesthetic State in Non-Human Primates

**DOI:** 10.1101/2021.09.12.459958

**Authors:** Sourish Chakravarty, Jacob Donoghue, Ayan S. Waite, Meredith Mahnke, Indie C. Garwood, Earl K. Miller, Emery N. Brown

## Abstract

Continuous monitoring of electroencephalogram (EEG) recordings in humans under general anesthesia (GA) has demonstrated that changes in EEG dynamics induced by an anesthetic drug are reliably associated with the altered arousal states caused by the drug. This observation suggests that an intelligent, closed-loop anesthesia delivery (CLAD) system operating in real-time could track EEG dynamics and control the infusion rate of a programmable pump to precisely maintain unconsciousness. The United States FDA acknowledges the potential benefits of such automatic physiological closed-loop control devices for patient care. Bringing these devices into clinical practice requires establishing their feasibility in suitable animal models. Therefore, given the close neurophysiological proximity between human and non-human primates (NHPs), we address this problem by developing and validating a propofol CLAD system in rhesus macaques. Our CLAD system has three key components: (1) a data acquisition system that records cortical local field potentials (LFPs) from an NHP in real-time; (2) a computer executing our CLAD algorithm that takes in the LFP signals as input and outputs infusion rates; and (3) a computer-controlled infusion pump that administers intravenous propofol. Our CLAD system controls an empirically-determined LFP marker of unconsciousness (MOU) at a user-prescribed target value by updating every 20 seconds the propofol infusion rate based on real-time processing of the LFP signal. The MOU is the instantaneous power in the 20 to 30 Hz band of the LFP spectrogram. Every cycle (duration *≈* 20 sec), our CLAD algorithm updates the MOU estimate and uses a robust optimal control strategy to adjust the propofol infusion rate based on the instantaneous error. This error is computed as the difference between the current and the user-prescribed target MOU values. Using neural recordings from multiple NHP anesthesia sessions, we first established that our chosen MOU signal was strongly correlated with propofol-induced decreased spiking activity which itself has been shown earlier to be associated with the level of unconsciousness in NHPs. Then we designed robust optimal control strategies that used subject-specific pharmacokinetic-pharmacodynamic models describing the MOU dynamics due to propofol infusion rate changes. Finally, we achieved safe and efficient closed-loop control of level of unconsciousness in 9 CLAD experiments involving 2 NHPs and 2 different 125 min long target MOU profiles with three target MOU changes within a given experiment. Our CLAD system performs stably, accurately and robustly across a total of 1125 min of closed-loop control. The CLAD performance measures, represented as median (25th percentile, 75th percentile), are 3.13 % (2.62%, 3.53%) for inaccuracy, 0.54 %(-0.31%, 0.89%) for bias, -0.02%/min (-0.06%/min, 0.00%/min) for divergence, and 3% (2.49%, 3.59%) for wobble. These performance measures were comparable or superior to previously reported CLAD performance measures from clinical studies (conducted outside USA) as well as rodent-based studies. The key innovations here are: (1) a pre-clinical NHP model for CLAD development and testing, (2) a neuroscience-informed LFP-based MOU for CLAD, (3) parsimonious, pharmacology-informed models to describe MOU dynamics under propofol infusion in rhesus macaques, (4) a novel numerical testing framework for propofol CLAD that incorporates a principled optimal robust control strategy for titrating propofol, and finally (5) experimental findings demonstrating the feasibility of stable, accurate and robust CLAD in the NHP model. Our NHP-based CLAD framework provides a principled pre-clinical research platform that can form the foundation for future clinical studies.

## 1 Introduction

General anesthesia is a drug-induced reversible state consisting of antinociception, unconsciousness, amnesia and immobility while maintaining physiological stability. Each year more than 300 million people worldwide receive general anesthesia to undergo surgery[1]. Being continuously attentive to the state of a patient during surgery and making frequent management decisions to optimize a patient’s care is fundamental to the practice of anesthesiology. To ensure a minimal level of vigilance for all patients, the American Society of Anesthesiologists has established standards for basic monitoring. They require the continuous presence of a qualified anesthesia care provider and continuous monitoring of oxygenation, ventilation, circulation and body temperature [2]. The anesthesiologist scans the relevant physiological variables (oxygen delivery, oxygen saturation, respiratory rate, heart rate, blood pressure and body temperature) on monitor displays, infers the patient’s physiological state, and makes any needed management to maintain the patient’s physiological state. The scanning cycle time of an anesthesia provider is usually 3 to 10 minutes. However, it can be longer if the provider becomes distracted with other management tasks. Automating this repetitive task would benefit patient care because its continual performance during moderate or long surgeries can cause fatigue and increase the chance of human error.

The best task to automate is monitoring and controlling the patient’s level of unconsciousness. Although anesthetic agents produce their primary effects in the brain and central nervous system, anesthesiologists are not required to monitor the brain state of a patient receiving anesthesia care. In fact, most do not. To guard against awareness, in the absence of brain monitoring, anesthesiologists most certainly administer anesthetic doses beyond what is needed to maintain an adequate level of unconsciousness. As a consequence, it is no surprise that brain dysfunction following general anesthesia is highly prevalent, particularly among the elderly [3].

A patient’s level of unconsciousness under general anesthesia can be reliably tracked using real-time processing of electroencephalogram (EEG) recordings [4, 5, 6, 7, 8]. This suggests that the EEG can be used to develop a closed-loop anesthesia delivery (CLAD) system to maintain precisely a specified level of unconsciousness. Since the 1950’s, EEG-based CLAD systems have been an active area of research [5, 9, 10, 11, 12, 13, 14, 15]. CLAD systems have been studied in rodents [16, 17], and outside the United States, in humans [10, 11, 12, 13, 14, 15]. The United States Food and Drug Administration (FDA) readily acknowledges the enormous benefits that CLAD systems, and more generally, Physiological Closed-Loop Control (PCLC) systems, can bring to patient care [18]. However, this regulatory body has approved no CLAD system for human testing in the United States. The FDA has identified testing of CLAD systems in appropriate large animal models as a critical prerequisite for ensuring performance reliability and reproducibility prior to human testing.

To address this critical gap, we develop and test a non-human primate (NHP) CLAD system for maintenance of unconsciousness using the anesthetic propofol. The system consists of: a data acquisition system that records neural signals in real-time from the cortex of an NHP, a computer that processes the neural signals using our CLAD algorithm and suggests an updated infusion rate at every 20 second interval, and a programmable infusion pump that administers propofol intravenously at the infusion rate suggested by the CLAD algorithm. The automatic titration is achieved by a feedback control framework that is designed for each NHP using an NHP-specific linear pharmacokinetics model (that characterizes the relationship between propofol infusion and its perceived effect in the brain) and a sigmoidal pharmacodynamics model (that characterizes the relationship between propofol’s brain concentration and an empirically-derived neurophysiological marker of unconsciousness (MOU)). We demonstrate our CLAD system with *in silico* testing as well as experimental testing in 2 NHPs during multiple experimental sessions.

## 2 Results

### 2.1 Developing a Propofol CLAD System for NHP Testing

We have developed a first instantiation of a propofol CLAD system and tested it in an NHP model (Fig. 1). In the current system, propofol is administered to an NHP as an intravenous infusion through an ear vein cannula using a computer-controlled syringe pump (Harvard Apparatus PhD Ultra). The LFP is recorded (Blackrock Cerebus neural data acquisition system [19]) from a Utah microelectrode array (Blackrock Microsystems [19]) implanted in the pre-frontal cortex. The user places the CLAD system in operation by setting a target marker of unconsciousness (MOU) level for the CLAD algorithm. The MOU is an empirically-defined function of the LFP that tracks level of unconsciousness. We establish that a reliable MOU for propofol in macaques is the normalized power in the 20 to 30 Hz band of the LFP spectrogram (Sec. 2.2). The CLAD algorithm controls the level of unconsciousness by adjusting the propofol infusion rate to control the MOU. The CLAD algorithm operates cyclically using two components: an MOU estimation framework and a control framework. The CLAD algorithm runs in a Matlab environment on a desktop computer and estimates in real time the current MOU level from the last 20 seconds of LFP data. Based on the the difference between target MOU level prescribed by the user and the estimated MOU level, the CLAD algorithm computes a new propofol infusion rate every 20 seconds.

**Figure 1:**
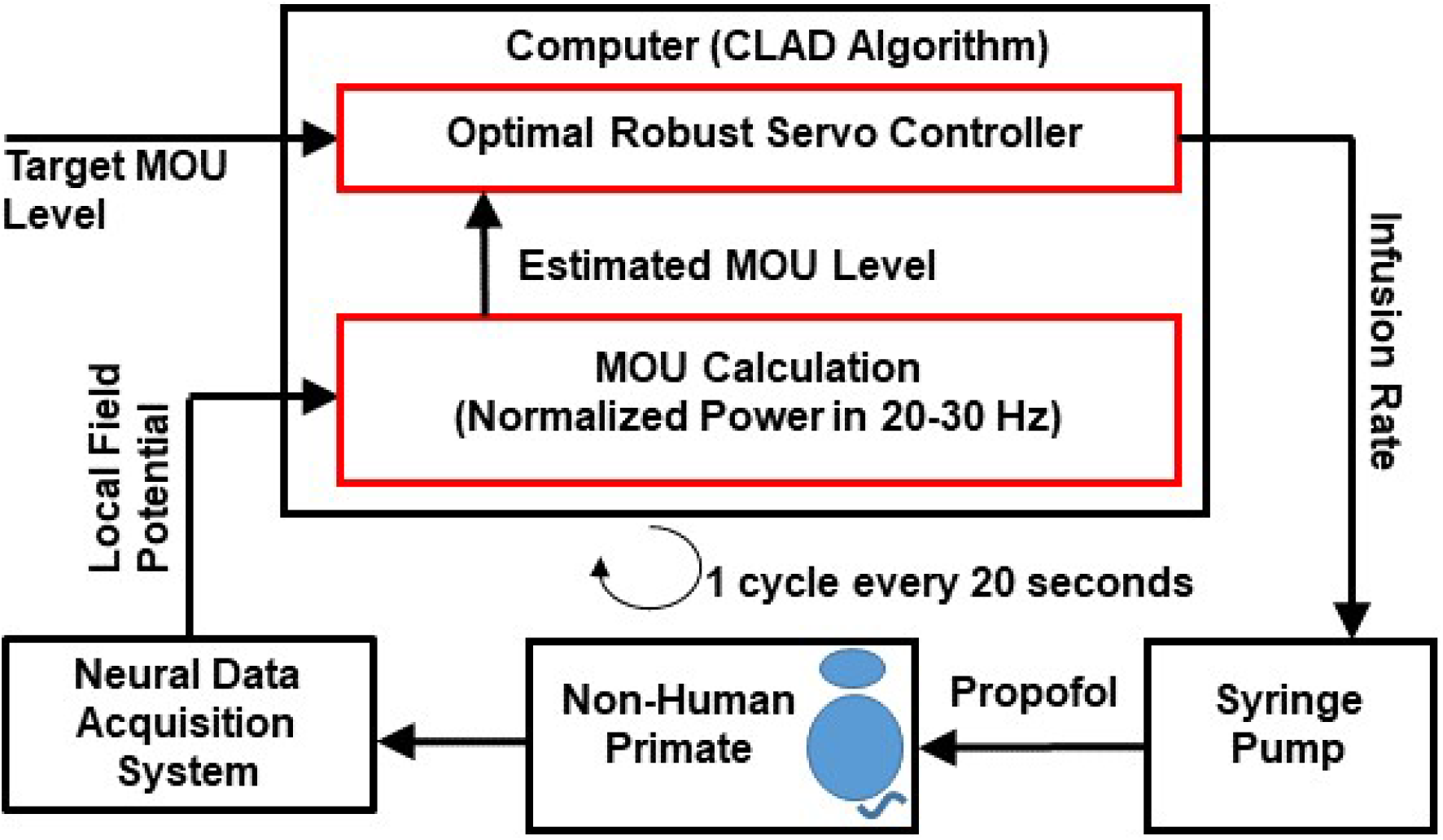
Schematic of our propofol-based CLAD in a NHP.

The control framework has three components: a linear two-compartment pharmacokinetics (PK) model (Fig. 3(A), upper panel), a sigmoidal-shaped pharmacodynamics (PD) model (Fig. 3(A), lower panel), and an optimal robust servo-control strategy [20, 21, 22]. The PK model describes the kinetics of how propofol reaches its effect-site, which is the brain. The PD model relates the propofol effect-site estimate to the MOU level. Together, the PK-PD model provide a subject-specific mathematical description of how the propofol infusion rate changes the MOU. The optimal robust servo-control strategy provides a principled way of leveraging the aforementioned PK-PD model to adjust the propofol infusion rate so as to achieve the user-prescribed target MOU level (see Sec. 4.5 for further elaboration).

Prior work has established NHP experimental models for studying the neurophysiological effects of propofol [23, 24] and other anesthetics [25, 26] on brain circuits. Using the experimental model reported by Bastos and colleagues [24], we recorded neural spiking activity and LFPs simultaneously while the animal received propofol administered by a computer-controlled infusion pump. The infusion rates were chosen to allow the animal to make the transition from the awake state to unconsciousness and back to the awake state. We analyzed the recordings to define the MOU, estimate PK and PD model parameters, and to determine the parameters for the optimal robust servo-controller.

### 2.2 The Baseline Normalized power in 20 to 30 Hz Range Was Chosen As The MOU to Control

To develop an MOU for use as a control variable in our propofol CLAD system, we studied the relationship between several candidate LFP-based spectral markers and neural spike rate. In the Methods (Sec. A) we provide details of the the NHP experimental setup (Sec. 4.1), spectral analysis (Sec. 4.2) and spike rate calculation (Sec. 4.3). Bastos et al. and Ishizawa et al. demonstrated that a decline in neural spike rate is a reliable indicator of unconsciousness maintained by propofol [24, 23]. Therefore, we analyzed the relationship between several candidate spectral markers and the neural spike rate as a function of time. Because we performed these analyses retrospectively, we had the profile of the entire infusion rate and behavioral events. An example of the data analyses from a single experimental session in NHP-B is presented in Fig. 2. In this experiment, propofol was administered at a constant infusion rate of 0.285 mg/kg/min for 60 minutes (Fig. 2A). Following Bastos and colleagues [24], we identified loss of consciousness as the time-point at which the animal closed its eyes and they remained closed for at least 5 minutes (Fig. 2C, eyes closed (EC), the first magenta vertical line). We identified recovery of consciousness as the time-point at which the animal opened its eyes for the first time after EC. (Fig. 2C, eyes open (EO), the second magenta vertical line). We defined the period of unconsciousness as the time between EC and EO (Fig. 2C). Of all the markers we considered, we identified the normalized total power in the 20 to 30 Hz band (Fig. 2B, 2C) as the one that tracked most faithfully changes in neural spike rate during the period of unconsciousness. Indeed, during unconsciousness, the Spearman’s rank correlation coefficient between the time course of the normalized power in the 20 to 30 Hz band and the time course of the normalized neural spike rate was 0.8760 (95% confidence interval [ 0.8729; 0.8791]) based on 11 recording sessions (Fig. 2D). Therefore, we took this spectral measure to be the propofol MOU. We normalized the MOU and the spike rate relative to the corresponding median value across the 5-minute period immediately prior to the start of propofol administration. Plots for all 11 recording sessions are in the Supplementary information (Figs. 6 and 7).).

**Figure 2:**
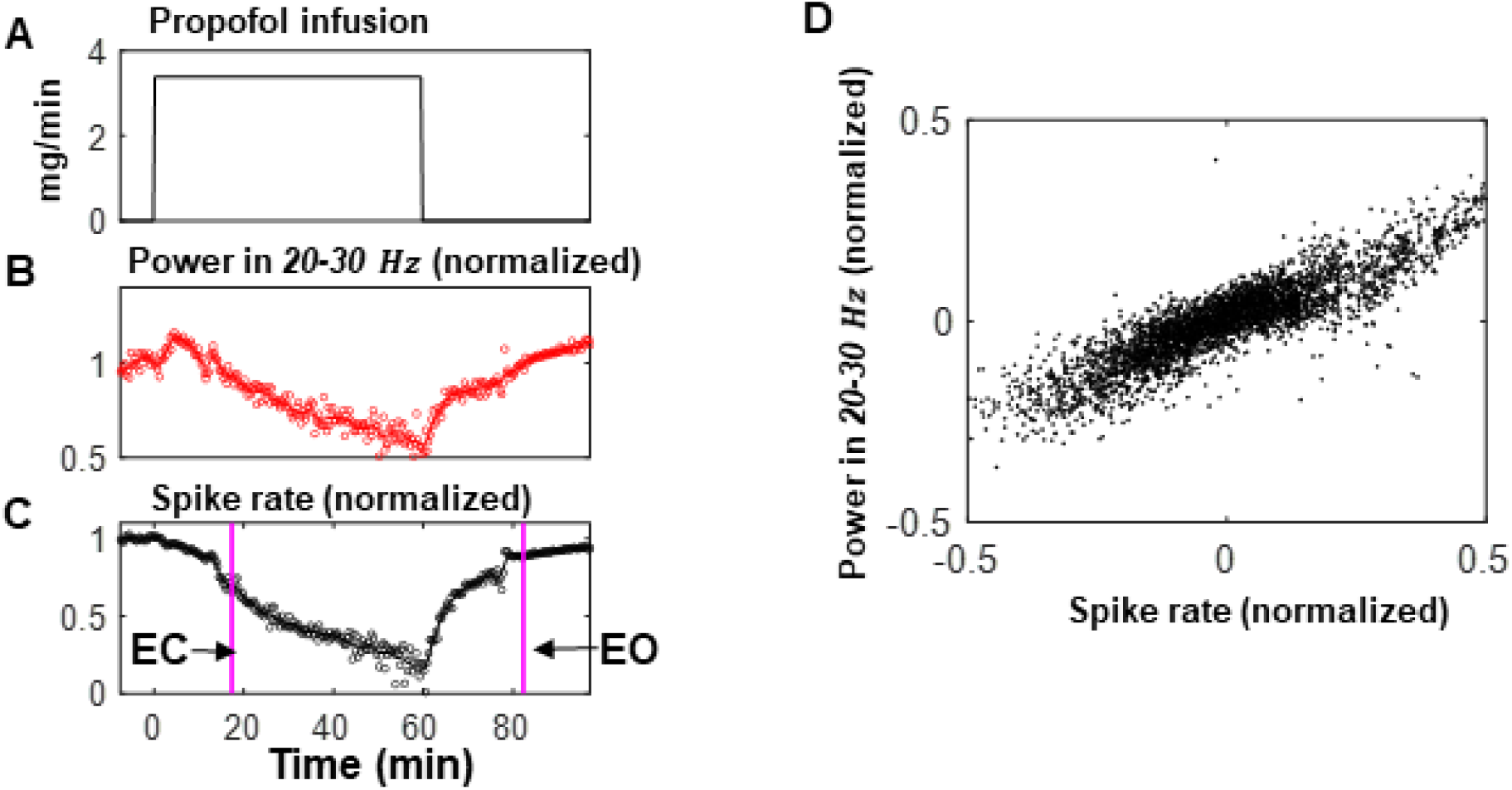
Neurophysiological response under propofol dosing in NHPs. Panels (A), (B), and (C) represent a single anesthesia session in NHP-B. (A) Infusion rate (B) baseline normalized LFP power in 20 – 30 Hz (bottom). (C) Baseline normalized spike rate. EC: Eyes Closed. EO: Eyes Open. (D) Plot of normalized power in 20-30 Hz vs. normalized spike rate across 11 anesthesia sessions in 2 monkeys (10 sessions in NHP-A). For consistency in representation across multiple recording sessions, the data from each recording session is centered about its respective mean value.

### 2.3 A Parsimonious Pharmacokinetic-Pharmacodynamic Model Reliably Describes Propofol’s MOU dynamics in NHP

We chose a linear two-compartment PK model (Fig. 3A, upper panel) because it is one of the simplest linear differential equation models that captures the essential features of how intravenous administration of propofol reaches the brain. The anesthetic enters the circulation or central compartment and passes to an inner compartment or the effect site, which we assume to be the brain and central nervous system [16, 17]. This model captures the essential feature that the MOU response to propofol changes at a slower rate or is delayed relative to the rate at which the anesthetic enters the circulation.

**Figure 3:**
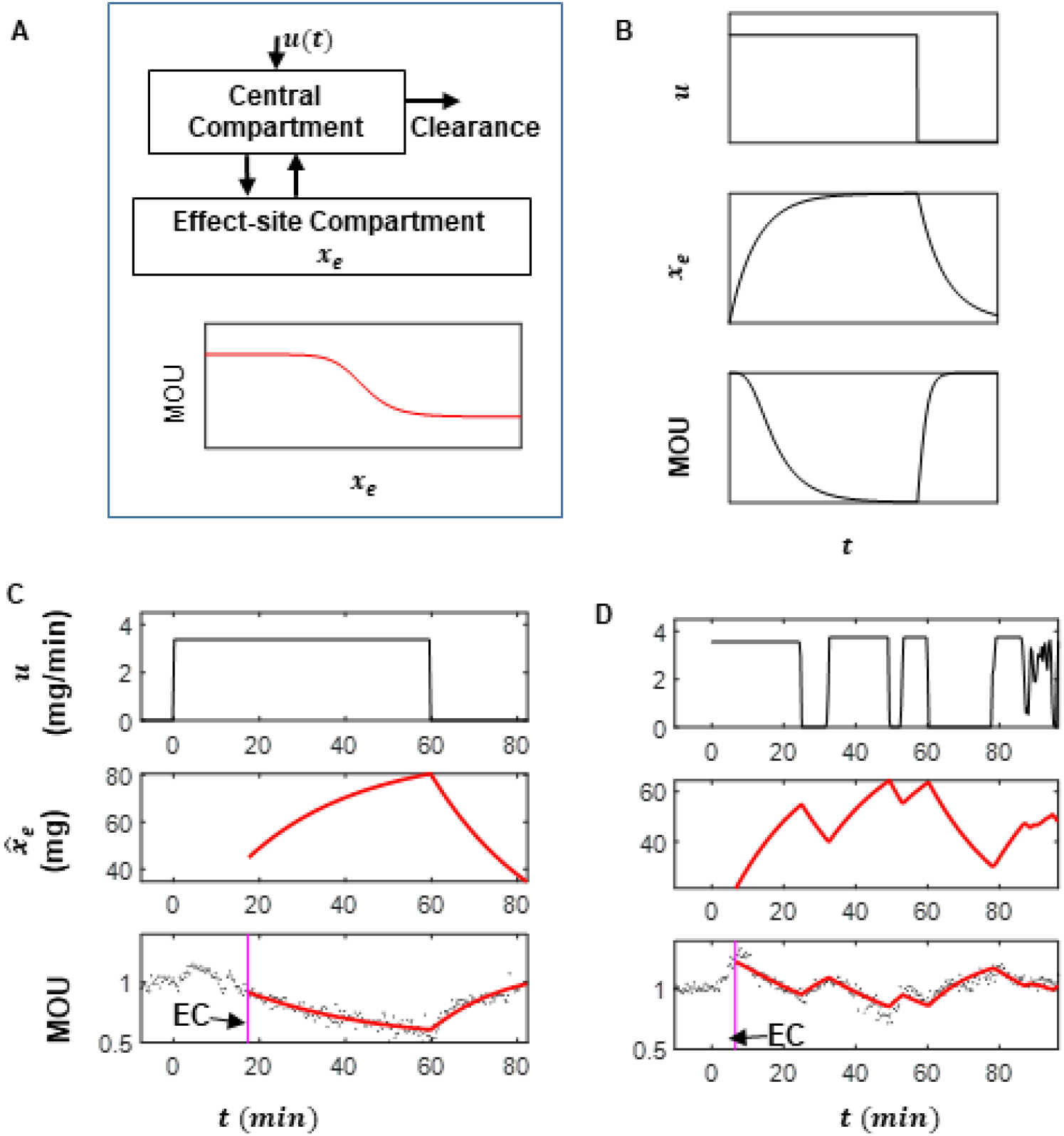
PK-PD model describing the propofol dose-dependent MOU dynamics, and PK-PD model fitting to dose response (see Sec. 4.4 for model description and estimation approach). A. Schematic of a two-compartment model, and a sigmoid model describing the dynamics in the MOU in terms of propofol effect-site level, *x*_*e*_, B. Simulated example of propofol infusion rate *u*, and corresponding effect-site level *x*_*e*_, and MOU trajectories. C and D represents PK-PD model fitting to a recording sessions from NHP-B (C) and another from NHP-A. (C, D; Top-panel) Stepped infusion protocol u, (C, D; middle panel) Estimated effect-site propofol level *x*_*e*_ (C, D; lower panel) Observed MOU (black dots) and MOU predicted by the best-fitting PK-PD model (solid red line). EC: Eyes Closed. The estimated parameters corresponding to sub-figure C are: *λ*_1_ = −0.0390, *λ*_2_ = −509.4326, *E*_0_ = 1.2009, *E*_*max*_ = 5.5551, *v*_50_ = 365.9014, and *γ* = 1.5118, with percentage residual error of 5.94%. The estimated parameters corresponding to sub-figure D are: *λ*_1_ = −0.0445, *λ*_2_ = −509.4326, *E*_0_ = 1.2807, *E*_*max*_ = 0.9299, *v*_50_ = 77.9048, and *γ* = 2.3845, with percent residual error of 5.13%.

For the PD model, we used a sigmoidal function to relate the MOU level and the propofol effect-site level (Fig. 3A, lower panel). We postulated based on prior anesthesia PD modeling studies [27, 28, 29] three phases in the neurophysiological response to propofol: a low response phase at low effect-site levels; a monotonically decreasing response phase in a middle range of effect-sites; and a second low response rate at high effect-site levels. The CLAD operation will ideally take place in the second phase where the MOU is most sensitive to changes in the propofol infusion rates. A simulated response of this PK-PD model under prolonged constant infusion is shown in Fig. 3B. For a constant propofol infusion rate the PK-PD model (Fig. 3B, top panel) produces a gradual increase in the effect-site level that approaches a steady-state value (Fig. 3B, middle panel). Assuming a sigmoidal relationship between the effect-site propofol level and the MOU, the MOU should mirror inversely the change in the effect-site level (Fig. 3B, middle and bottom panels).

We fit the combined PK-PD model (Fig. 3A) to the MOU versus time and infusion rate versus time data from the period of unconsciousness for a recording session from NHP-A and one from NHP-B (Fig. 3C, 3D). For the PK-PD model fitting procedure we used a cyclic descent non-linear least squares optimization approach [30] to estimate simultaneously the PK and the PD model parameters, and the time courses of the propofol effect-site levels (see Sec. 4.4 for details). The time course of the estimated propofol effect-site level (Fig. 3C, 3D, middle panel) is consistent with the time course of the infusion profile (Fig. 3C, 3D), top panel). In Fig. 3, the recording session from NHP-B had a constant infusion profile whereas the session from NHP-A had a variable infusion profile, both with a maximum infusion rate of 0.4 mg/kg/min. The analysis corroborates the inverse relationship between the MOU (Fig. 3C, bottom panel) and the estimated effect-site propofol level (Fig. 3C middle panel) as implied by our model construction (Fig. 3B, middle and bottom panels). The periods of unconsciousness used for data fitting were 65 minutes for NHP-B and 90 minutes for NHP-A. For both the animals the 5-parameter PK-PD model provided a reasonable description of the pharmacodynamic response of the MOU. The PK-PD models identified from these two sessions were subsequently used to develop NHP-specific control system for the CLAD.

### 2.4 An Optimal Robust Servo-controller Can Reliably Control MOU in Computer Simulations

For our control strategy we combined the Linear Quadratic Regulator (LQR) strategy with estimated state feedback used in our previous rodent CLAD studies [16, 17], and the PID control framework used in human CLAD studies outside the USA [31, 32, 15]. We verified in a simulation study that this hybrid approach, with the principled optimization-based design framework of the former and the stable output tracking property under model uncertainty of the latter, could stably control the pharmacodynamic response of propofol PK-PD models of human subjects [33]. This inspired us to choose as our feedback control strategy an optimal robust servo-control scheme that updates the infusion rate by combining two components: an LQR strategy and an integral compensation strategy that monitors accumulation of the tracking error (*MOU*_*target*_ − *MOU*_*actual*_) [21, 20, 22]. A detailed block diagram of the feedback control strategy is presented in Supplementary Fig. 8; Fig. 4A provides a simplified illustration of the same.

**Figure 4:**
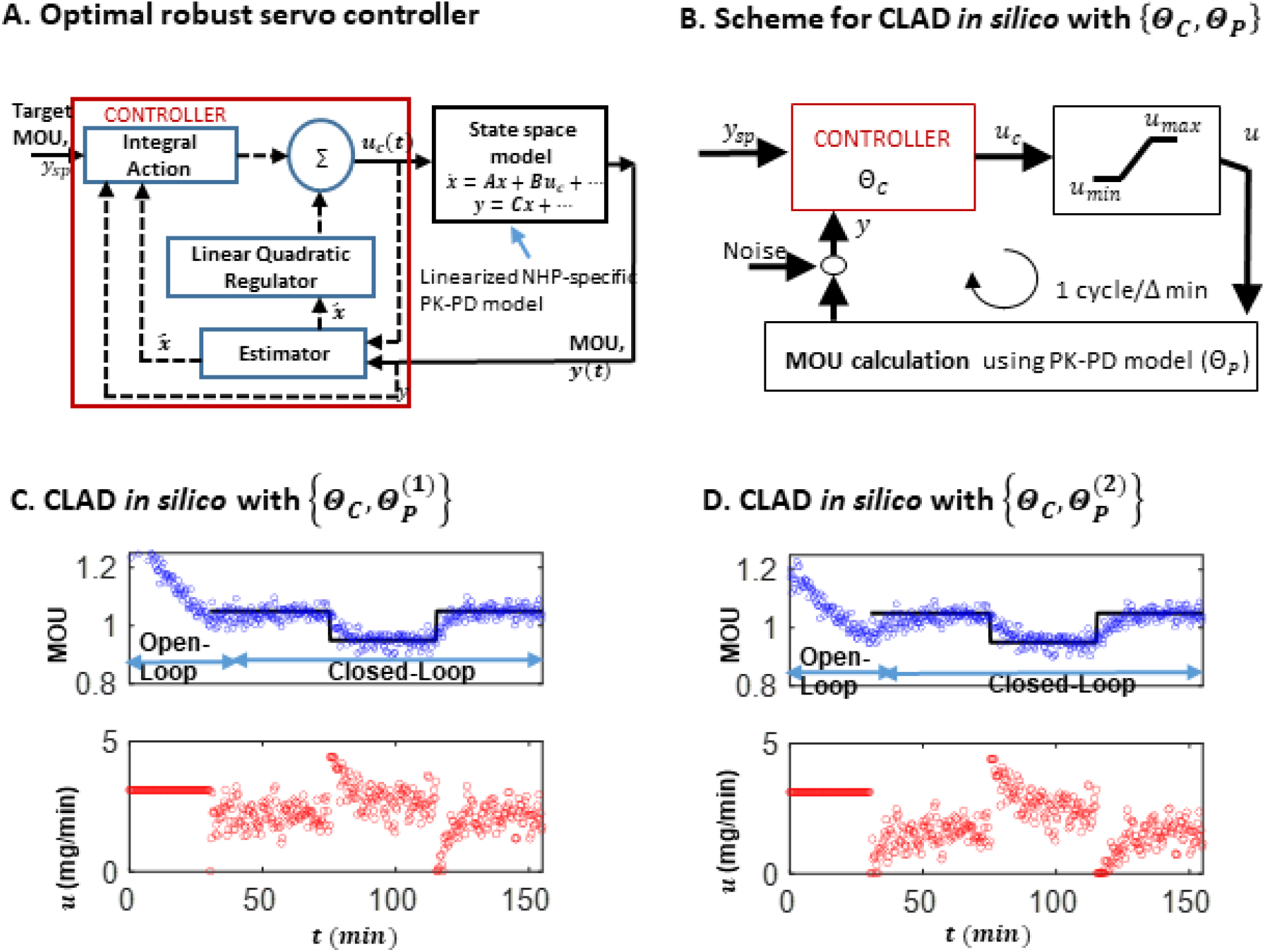
Block diagram of *in silico* control simulation, and simulation results.(A) Schematic block diagram of the linear optimal robust servo controller, (B) Schematic of *in silico* testing of CLAD algorithm comprising a linear controller (set of parameters characterizing the controller design is denoted by Θ_*C*_), a control saturation block to explicitly bound the suggested infusion rate, *u*_*c*_ by the linear controller in the range [*u*_*min*_, *u*_*max*_], and a PK-PD model (set of parameters characterizing a PK-PD model is denoted by Θ_*P*_). (C, D) Two simulated CLAD sessions where the top panel shows Target MOU levels (horizontal black line), and estimated MOU level (blue circles), and the bottom panel presents Infusion rates (red circles). The controller parameters are same (Θ_*C*_) in both sessions, but the system parameters are different. This is indicated by 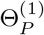 and 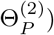) denoting the the PK-PD model used for MOU calculation in C and D, respectively. Details on controller design are provided in Methods (sec. 4.5).

The LQR component uses a state estimator to compute the propofol effect-site level [34]. Since the LQR coupled to an observer can be sensitive to model misspecification, we included an integral compensation that can ensure stable MOU tracking even when the PK-PD model is misspecified. Hence, the integral compensation helps ensure a degree of robustness in our control strategy. In addition, the controller framework provides a principled optimization-based paradigm to design a two-degree controller for a linear system with step target MOU values and step disturbances. The two-degree of freedom feature of the controller stems from the fact that the response to step target MOU values and to step disturbances can be tuned separately without affecting the performance of either [21, 20, 22]. For the optimal robust servo-control this is done by minimizing two distinct quadratic criteria. The solution of the first leads to the LQR feedback gains and the solution to the second characterizes the integral action and state estimator gains (see [21, 20, 22] for details). The control design is characterized by the set of user-prescribed parameters that goes in the definitions of the quadratic criteria.

Choosing the feedback control strategy is just the first step towards our CLAD controller design. The next step is tuning the parameters of the controller so that the CLAD system has desired properties. Since the chosen feedback control strategy was developed for linear systems [21, 20, 22], we linearized the NHP PK-PD model about a steady state infusion rate of 0.285 mg/kg/min - a value that we determined empirically to be a safe dose for maintaining unconsciousness during a prolonged experimental session for an NHP [24]. We then manually chose parameters that characterized the two aforementioned quadratic criteria so that the linearized closed-loop system had high phase margin (*>* 80 degree), high gain margin (*>* 60 dB) and low settling times for a unit step input (*<* 11 min). See Methods (sec. 4.5) for a brief explanation of these design criteria.

Using the scheme in Fig. 1, we tested *in silico* the ability of this linear feedback control strategy to provide acceptable tracking performance when the dose-effect site relationship was governed by the NHP’s PK-PD model. To mimic actual experimental conditions in our simulations, we set the lower (upper) infusion rate limit at 0.0 (0.4) mg/kg/min, added Gaussian white noise to the PK-PD model output, and enforced fixed rate of infusion between each discrete controller updates. The controller updates were carried out every 20 seconds. We found that the control framework using the same single set of manually tuned parameters worked well with multiple PK-PD model parameter sets. The inaccuracy was less than 5% across these simulations.

Since MOU values around 1 corresponded to appropriate levels of unconsciousness post-EC in the prior open-loop experimental sessions (Fig. 6), the target MOUs for both *in silico* and experimental CLAD studies were modulated around 1. An MOU value less than (greater than) 1 would indicate that the animal is at a more profound (less profound) level of unconsciousness. We tested our propofol CLAD system (Fig. 1) in a 155-minute in silico experiment (see Fig. 4C) divided into 4 stages. During the first 30 minutes (stage 1), the system is run in an open loop mode by setting constant propofol infusion rate of 0.285 mg/kg/min. During the remaining 125 minutes of the experiment, the CLAD system controlled the propofol infusion rate for 45 minutes to maintain the MOU at 1.05 (stage 2), next for 40 minutes to maintain the MOU at 0.95 (stage 3), and finally for a final 40 minutes at 1.05 again (stage 4). In the simulation framework for this experiment (Fig. 4B) the PK-PD model parameter (in the MOU calculation block) is set to the estimated PK-PD model (from NHP-A) that was linearized to develop the linear feedback controller (Fig. 4A).

To simulate the scenario typically encountered in the operating room, the experiment began with the propofol infusion rate set at the constant value of 0.285 mg/kg/min (see Fig. 4C) for the first 30 minutes. Not surprisingly, with a constant infusion rate the MOU does not remain constant but instead, monotonically decreases suggesting that the animal is moving to a more profound level of unconsciousness (Fig. 4C, top-panel, blue circles). After the initial 30-minute period, the CLAD system takes over control of the propofol infusion rate for the first 45 minutes of closed-loop control with a target MOU value of 1.05 (stage 2). To decrease the level of unconsciousness to the target of 1.05 (Fig. 4C, top panel, solid horizontal black line), the CLAD system stops the infusion for 5 minutes (Fig. 4C, bottom panel). Once the target level of 1.05 was acquired, the infusion restarted and was maintained approximately around a mean infusion rate of 0.20 mg/kg/min. During the next 40 minutes of closed-loop control (stage 3), the target MOU was set at the lower value of 0.95 meaning a more profound level of unconsciousness. The CLAD system achieves the new target within approximately 8 minutes (Fig. 4C, top panel) by initially increasing the infusion rate to its maximum value of 0.4 mg/kg/min then maintaining the rate at an average of approximately 0.25 mg/kg/min (Fig. 4C, bottom panel). For the final 40 minutes of closed-loop control (stage 4), the target level was again set at 1.05 (Fig. 4C, top panel). The CLAD system again achieves the new target level by turning the infusion off. Once the target level of 1.05 was achieved, the CLAD system controlled the MOU maintaining an average infusion rate of approximately 0.20 mg/kg/min.

Fluctuation around this mean due to the CLAD system’s 20-second updates are critical for maintaining the MOU at the constant value of 1.05. The overall inaccuracy in the controller’s target tracking was only 1.64% for 125 min of closed-loop control. The other control performance metrics, bias, wobble, and divergence were respectively -0.50%, 1.62% and -0.00 %/min (definitions and interpretations of CLAD performance measures are provided in Sec 4.6). These findings establish the feasibility of using the optimal robust servo controller (designed for a linearized PK-PD model ; Fig. 4A) along with the input saturation block (Fig. 4B) to stably control a user-prescribed MOU.

To further verify that the control design can reliably perform for a set of PK-PD parameters different from those that were used for controller tuning, the PK-PD parameters in the MOU calculation block in Fig. 4B were set to values that were physically realizable, but different from the ones used for controller design (Fig. 4D). Even with this model misspecification, the target tracking in Fig. 4D was stable, and the time to target convergence and performance metrics (inaccuracy of 1.96%, bias of -0.76%, wobble of 1.86% and divergence of -0.01 %/min) were similar to those observed in Fig. 4C.

### 2.5 An Optimal Robust Servo-controller Reliably Controls The Target MOU in NHP Experiments

We tested the propofol CLAD system *in vivo* (Fig. 5A-I) in 9 experimental sessions (each 155-minute long) that followed the structure of the *in silico* experiments. In the first 30 minutes the CLAD system operated in an open-loop mode with a constant propofol infusion rate. During the remaining 125 minutes, the CLAD system controlled the propofol infusion rate for 45 minutes with the MOU at 1.05, 40 minutes at 0.95 and the final 40 minutes again at 1.05. To again emulate, the typical practice in the operating room, we start the experiment with the propofol infusion at the constant rate of 0.285 mg/kg/min (see bottom panel of any of the sub-figures Fig. 5A) for the first 30 minutes. This rate was again intended to achieve an MOU level around 1 as in the *in silico* experiment. As in *in silico* experiments (Fig. 4C and 4D), when the infusion rate was held constant, the MOU decreased monotonically indicating that the animal was moving to a more profound level of unconsciousness (blue circles in the top panel of any of the sub-figures Fig. 5A).

**Figure 5:**
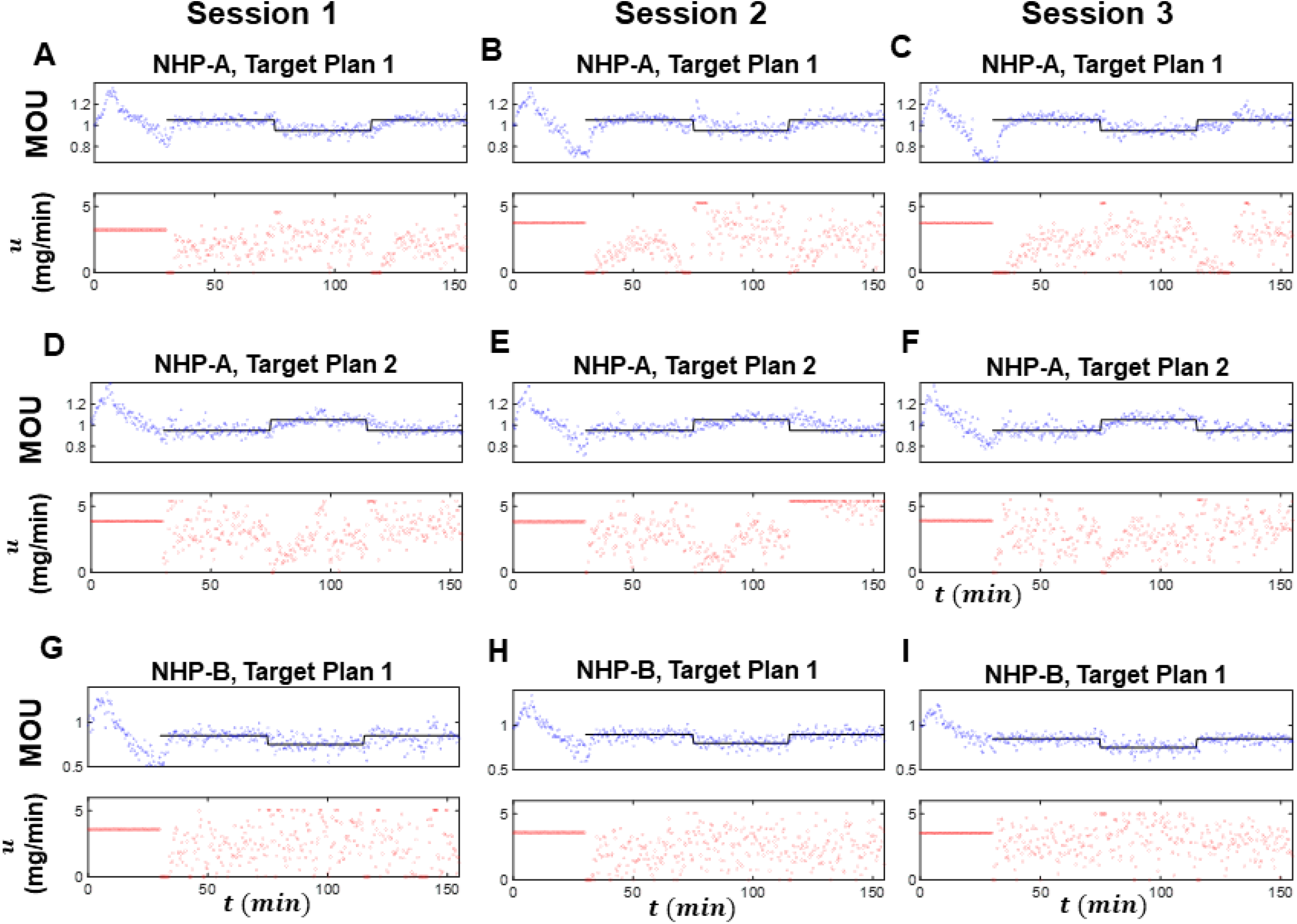
Experimental validation session in two NHPs. Sub-figures A, B, and C present the experimental observation from 3 CLAD sessions in NHP-A with target set to an inverted top-hat profile (Target Plan 1) after a 30 min of constant infusion of propofol. Sub-figures D, E, and F present data from another 3 CLAD sessions from NHP-A with similar design as A, B or C, but with the target set to a top-hat profile (Target Plan 2). Sub-figures G, H and I present data from 3 CLAD sessions from NHP-B with Target Plan 1.

At minute 30, the CLAD system took over control of the propofol infusion rate to initiate the first 45 minute of closedloop control. Again, the CLAD system updated the propofol infusion rate every 20 seconds (approximately 20 seconds to be precise, since there can be an additional delay of approximately 1 second stemming from the real-time data transfer and computing involved in an iteration of the CLAD algorithm). Similar to the *in silico* experiment, to achieve the decreased the level of unconsciousness, 1.05 (Fig. 5A, upper panel, solid black line), the CLAD system turned off the infusion for approximately 3 minutes (Fig. 5A, lower panel). Once the target level of 1.05 was acquired, the infusion restarted, and the rate fluctuates appreciably from update to update to keep the MOU well maintained at the target level. During the second 40 minutes of closed-loop control, the target was decreased to the lower marker value of 0.95, meaning a deeper level of unconsciousness. The CLAD system achieved the new target within approximately 4 minutes (Fig. 5A, top panel). The infusion rate fluctuated appreciably to control the MOU (Fig. 5A, bottom panel). For the final 40 minutes of closed-loop control the target level was reset to 1.05 (Fig. 5A). The CLAD system again achieved the new target level by turning the infusion off for approximately 4 minutes. Once the target level of 1.05 was achieved, the CLAD system’s infusion rates fluctuated appreciably to maintain tight control of the MOU.

To determine if performance in the same subject could be replicated across multiple sessions with the same target plan, we repeated the experiment in Fig. 5A two more times (Figs. 5B and C). The performance metrics from the 3 sessions are in Table 1(Expt. 1-3). To analyze whether the performance was robust across different target profiles, We performed three CLAD experiments with NHP-A and a protocol identical to that in Fig. 5A-C, with the exception that the MOU target trajectory was changed from an inverted top-hat profile to a top-hat profile. After 30 minutes of a constant infusion at 0.285 mg/kg/min, we initiated the top-hat protocol. It consists of 45 minutes of closed-loop MOU control at level 0.95, 40 minutes of closed-loop MOU control at level 1.05; and 40 minutes of closed-loop MOU control at level 0.95. The CLAD performance metrics across these sessions are tabulated in Table 1 (Expt. 4-6). To further analyze the robustness of the CLAD performance to subject-wise variability, 3 CLAD experiments were performed in NHP-B with a near-identical protocol to the one in Fig. 5(A-C). We retained the inverted top-hat profile and a difference of 0.1 between consecutive target levels. The CLAD performance metrics across these sessions are tabulated in Table 1 (Expt. 7-9). Note that the exact MOU target levels in NHP-B (Expts. 7-9 in Table 1) was adjusted while maintaining the same inverted top-hat profile of target plan 1 in NHP-A as the MOU target levels prescribed for NHP-A (Expts. 1-6 in Table 1) were not appropriate to maintain the desired level of unconsciousness in NHP-B. Due to a similar observation between sessions in NHP-B, the exact target MOU levels in NHP-B were further adjusted (compare target MOU values in Expt. 7 - 9 in Table 1).

**Table 1:** CLAD performance measures (see Sec. 4.6) from 3 independent repeat sessions of CLAD: in NHP-A with Target Plan 1 (inverted top-hat profile) (Expt.#: 1, 2, 3), in NHP-A with Target Plan 2 (top-hat profile) (Expt.#: 4, 5, 6), and in NHP-B with Target Plan 1 (Expt. #: 7, 8, 9). For each experimental session, the data segments, L1, L2, and L3 comprise the last 30 min of data in each target level of the prescribed target plan before a target level switch occurs. The time intervals for L1, L2 and L3 are therefore, respectively, [45 min, 75 min], [85 min, 115 min], [125 min, 155 min], where the time point 0 min marks the beginning of the initial open-loop infusion mode. The median and the 25th and 75th percentiles are reported.

| Expt.# | NHP/Target Plan/Rep.<br>/Segment (target MOU) | MDAPE<br>(%) | MDPE<br>(%) | Div<br>(%/min) | Wob<br>(%) | $u_{avg}$<br>(mg/kg/min) | $u_{std}$<br>(mg/kg/min) |
| --- | --- | --- | --- | --- | --- | --- | --- |
| 1 | NHP-A/1/1/L1 (1.05) | 2.30 | -0.28 | 0.06 | 2.34 | 0.17 | 0.07 |
|  | NHP-A/1/1/L2 (0.95) | 3.54 | 0.54 | -0.01 | 3.60 | 0.23 | 0.09 |
|  | NHP-A/1/1/L3 (1.05) | 2.37 | 0.89 | 0.03 | 2.35 | 0.20 | 0.06 |
|  | Median | 2.37 | 0.54 | 0.03 | 2.35 | 0.20 | 0.07 |
|  | (25th, 75th) | (2.32, 3.25) | (-0.07, 0.80) | (-0.00, 0.05) | (2.34, 3.29) | (0.18, 0.22) | (0.07, 0.09) |
| 2 | NHP-A/1/2/L1 (1.05) | 2.35 | -0.13 | 0.12 | 2.47 | 0.12 | 0.07 |
|  | NHP-A/1/2/L2 (0.95) | 3.44 | 0.57 | -0.01 | 3.53 | 0.24 | 0.08 |
|  | NHP-A/1/2/L3 (1.05) | 3.16 | 0.32 | 0.04 | 2.99 | 0.19 | 0.08 |
|  | Median | 3.16 | 0.32 | 0.04 | 2.99 | 0.19 | 0.08 |
|  | (25th, 75th) | (2.55, 3.37) | (-0.01, 0.51) | (0.00, 0.10) | (2.60, 3.39) | (0.14, 0.23) | (0.07, 0.08) |
| 3 | NHP-A/1/3/L1 (1.05) | 2.74 | 1.80 | -0.04 | 2.22 | 0.16 | 0.06 |
|  | NHP-A/1/3/L2 (0.95) | 2.61 | 0.67 | -0.00 | 2.54 | 0.22 | 0.07 |
|  | NHP-A/1/3/L3 (1.05) | 3.24 | 1.80 | -0.16 | 2.96 | 0.20 | 0.11 |
|  | Median | 2.74 | 1.80 | -0.04 | 2.54 | 0.20 | 0.07 |
|  | (25th, 75th) | (2.65, 3.11) | (0.95, 1.80) | (-0.13, -0.01) | (2.30, 2.86) | (0.17, 0.21) | (0.06, 0.10) |
|  | Median (Expt.#1-3)<br>(25th, 75th) | 2.74<br>(2.37, 3.29) | 0.57<br>(0.21, 1.12) | -0.00<br>(-0.02, 0.05) | 2.54<br>(2.34, 3.12) | 0.20<br>(0.17, 0.22) | 0.07<br>(0.07, 0.09) |
| 4 | NHP-A/2/1/L1 (0.95) | 2.88 | -0.63 | -0.08 | 3.00 | 0.23 | 0.07 |
|  | NHP-A/2/1/L2 (1.05) | 2.30 | -0.44 | -0.02 | 2.22 | 0.19 | 0.08 |
|  | NHP-A/2/1/L3 (0.95) | 2.62 | -0.04 | -0.02 | 2.58 | 0.26 | 0.06 |
|  | Median | 2.62 | -0.44 | -0.02 | 2.58 | 0.23 | 0.07 |
|  | (25th, 75th) | (2.38, 2.81) | (-0.58, -0.14) | (-0.06, -0.02) | (2.31, 2.89) | (0.20, 0.25) | (0.06, 0.08) |
| 5 | NHP-A/2/2/L1 (0.95) | 3.76 | 0.07 | -0.02 | 3.75 | 0.23 | 0.08 |
|  | NHP-A/2/2/L2 (1.05) | 2.63 | 0.56 | -0.06 | 2.47 | 0.18 | 0.08 |
|  | NHP-A/2/2/L3 (0.95) | 3.47 | 2.11 | -0.15 | 3.37 | 0.38 | 0.03 |
|  | Median | 3.47 | 0.56 | -0.06 | 3.37 | 0.23 | 0.08 |
|  | (25th, 75th) | (2.84, 3.68) | (0.19, 1.72) | (-0.13, -0.03) | (2.70, 3.66) | (0.19, 0.34) | (0.04, 0.08) |
| 6 | NHP-A/2/3/L1 (0.95) | 3.50 | -0.39 | -0.09 | 3.55 | 0.21 | 0.08 |
|  | NHP-A/2/3/L2 (1.05) | 2.57 | 1.01 | 0.00 | 2.38 | 0.19 | 0.07 |
|  | NHP-A/2/3/L3 (0.95) | 2.62 | 0.89 | -0.03 | 2.64 | 0.25 | 0.07 |
|  | Median | 2.62 | 0.89 | -0.03 | 2.64 | 0.21 | 0.07 |
|  | (25th, 75th) | (2.59, 3.28) | (-0.07, 0.98) | (-0.08, -0.00) | (2.45, 3.32) | (0.20, 0.24) | (0.07, 0.08) |
|  | Median (Expt.#4-6)<br>(25th, 75th) | 2.63<br>(2.61, 3.48) | 0.07<br>(-0.40, 0.92) | -0.03<br>(-0.08, -0.02) | 2.64<br>(2.45, 3.42) | 0.23<br>(0.19, 0.25) | 0.07<br>(0.06, 0.08) |
| 7 | NHP-B/1/1/L1 (0.85) | 4.08 | 0.63 | -0.03 | 3.96 | 0.17 | 0.10 |
|  | NHP-B/1/1/L2 (0.75) | 5.59 | 1.47 | -0.08 | 5.61 | 0.24 | 0.11 |
|  | NHP-B/1/1/L3 (0.85) | 6.81 | -2.02 | -0.05 | 6.60 | 0.16 | 0.13 |
|  | Median | 5.59 | 0.63 | -0.05 | 5.61 | 0.17 | 0.11 |
|  | (25th, 75th) | (4.46, 6.51) | (-1.36, 1.26) | (-0.07, -0.04) | (4.37, 6.35) | (0.16, 0.22) | (0.10, 0.13) |
| 8 | NHP-B/1/2/L1 (0.90) | 3.01 | -0.42 | 0.02 | 3.17 | 0.16 | 0.08 |
|  | NHP-B/1/2/L2 (0.80) | 4.25 | 1.02 | -0.06 | 4.12 | 0.21 | 0.10 |
|  | NHP-B/1/2/L3 (0.90) | 3.43 | 0.09 | -0.02 | 3.38 | 0.20 | 0.09 |
|  | Median | 3.43 | 0.09 | -0.02 | 3.38 | 0.20 | 0.09 |
|  | (25th, 75th) | (3.11, 4.05) | (-0.29, 0.78) | (-0.05, 0.01) | (3.22, 3.94) | (0.17, 0.21) | (0.09, 0.10) |
| 9 | NHP-B/1/3/L1 (0.85) | 3.13 | -0.60 | 0.00 | 2.82 | 0.21 | 0.08 |
|  | NHP-B/1/3/L2 (0.75) | 5.30 | 0.55 | -0.10 | 5.36 | 0.27 | 0.11 |
|  | NHP-B/1/3/L3 (0.85) | 3.04 | -0.32 | -0.02 | 3.21 | 0.22 | 0.08 |
|  | Median | 3.13 | -0.32 | -0.02 | 3.21 | 0.22 | 0.08 |
|  | (25th, 75th) | (3.06, 4.76) | (-0.53, 0.33) | (-0.08, -0.00) | (2.92, 4.82) | (0.22, 0.25) | (0.08, 0.10) |
|  | Median (Expt.#7-9)<br>(25th, 75th) | 4.08<br>(3.11, 5.38) | 0.09<br>(-0.46, 0.73) | -0.03<br>(-0.07, -0.01) | 3.96<br>(3.20, 5.42) | 0.21<br>(0.17, 0.22) | 0.10<br>(0.08, 0.11) |
|  | Median (Expt.#1-9)<br>(25th, 75th) | 3.13<br>(2.62, 3.53) | 0.54<br>(-0.31, 0.89) | -0.02<br>(-0.06, 0.00) | 3.00<br>(2.49, 3.59) | 0.21<br>(0.18, 0.23) | 0.08<br>(0.07, 0.09) |

The median (25th percentile, 75th percentile) of CLAD performance measures (MDAPE, MDPE, divergence, and wobble) as well as the median and standard deviation of the infusion rates across all 27 data segments (3 segments per experiment × 9 experiments) were: MDAPE of 3.13 % (2.62%, 3.53%), MDPE of 0.54 %(-0.31%, 0.89%), divergence of -0.02%/min (-0.06%/min, 0.00%/min), and wobble of 3% (2.49%, 3.59%). These values are comparable to earlier propofol-based CLAD studies in rodents by Shanechi et al. where MDAPE of 3.61% and MDPE of -1.44% were reported [16]). Also the performance measure values observed here are comparable to those reported in an early CLAD study by Struys et. al. (MDAPE of 7.7 ± 2.49, MPDE of −6.6 ± 2.63, divergence of 0.024 ± 0.029, wobble of 5.90 ± 2.33 per Struys et. al. [35]), and lower than the values reported in a more recent CLAD study by Puri et al. (median (25th percentile, 75th percentile) values of 10% (10%, 12%) for MDAPE and 9% (8%, 10%) for wobble per Puri et. al. [15]). The low values for the median of performance measures in the current work suggests that the accuracy of the CLAD system is acceptable based on the benchmarks set in earlier propofol-based CLAD studies [16, 35, 15]. The narrow inter-quartile ranges for these four metrics reported in our NHP studies suggests that the CLAD system performance was robust to intra-session, inter-session and inter-subject variability.

Using the performance measures reported in Table 1, we compare across three groups: NHP-A with target plan 1, NHP-A with target plan 2, and NHP-B with target plan 1. For each group and for a given performance measure, we assume the performance measure calculated from the nine 30 min segments (three 30 min segments per experiment × three repeats) as independent observations for that group. Box plots in supplementary Fig. 9 allows for visual comparison across these three groups. For a more precise comparison under additional assumption of normality of performance measure within each group and equal variance across the three groups and null hypothesis that the mean values are similar across these 3 groups, we perform one-way ANOVA followed by pair-wise comparison with Bonferroni correction (Table 2, 3, 4, 5, 6, 7). Based on the one-way ANOVA analysis, we infer that mean MDAPE is similar between NHP-A with target plan 1 and NHP-A with target plan 2 experiments, but both these groups have a lower MDAPE relative to NHP-B with target plan 2. This agrees with the visual inspection in Fig. 9A. The one-way ANOVA analysis suggests that the mean MDPE on the other hand is similar across these three groups. This agrees with visual inspection in Fig. 9B. Although visual inspection of Fig. 9C suggest that divergence is higher in NHP-A relative to NHP-B for the same target plan, the one-way ANOVA test is unable to reject the null hypothesis that the mean divergence is same across these groups. The one-way ANOVA analysis indicates that the mean wobble is similar between NHP-A with target plan 1 and NHP-A with target plan 2 experiments, but both these groups have a lower wobble relative to NHP-B with target plan 1. This agrees with the visual inspection of Fig. 9D. For the average infusion rate, both the ANOVA test and the visual inspection of Fig. 9E suggests that the average infusion rates are similar across the groups. The slightly higher value of *u*_*avg*_ observed in NHP-A with target plan 2 relative to NHP-A with target plan 1 can be explained by the fact that the former (with top-hat target profile) has a higher fraction of the experimental duration conducted at a lower MOU target value, or, deeper level of unconsciousness and thus requiring higher propofol exposure. The one-way ANOVA analysis indicates that the mean *u*_*std*_ is similar between NHP-A with target plan 1 and NHP-A with target plan 2 experiments, but both these groups have a lower mean *u*_*std*_ relative to NHP-B with target plan 2. This agrees with the visual inspection of Fig. 9F. The higher values of inaccuracy (characterized by MDAPE), intra-session time-related variation (characterized by wobble), and corresponding higher control fluctuations (characterized by *u*_*std*_) observed in NHP-B relative to NHP-A could plausibly be attributed to the difference in propofol-driven neurophysiogical response that was not sufficiently accounted in the controller design for NHP-B.

**Table 2:**
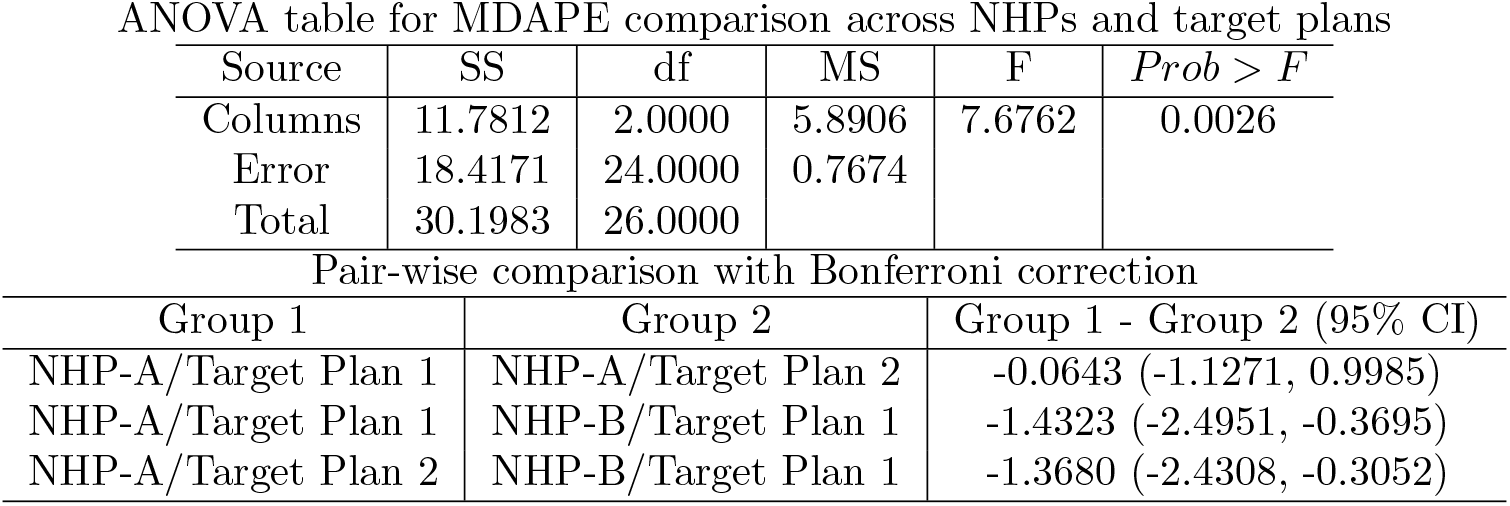
Comparing MDAPE across 2 NHPs and 2 target plans. One-way ANOVA table and pairwise comparison table reported. Each of the 3 groups comprised of 9 values of MDAPE = 3 per experiments × 3 repeat experiments. SS - sum of squares, df - degree of freedom, MS - mean squared errors, F-estimated F-statistic which is the ratio of the mean squared errors, *Prob > F* denotes the probability that the ratio of the mean squared errors exceeds the estimated F-statistic, CI - confidence interval

| Source | SS | df | MS | F | *Prob > F* |
| --- | --- | --- | --- | --- | --- |
| Columns | 11.7812 | 2.0000 | 5.8906 | 7.6762 | 0.0026 |
| Error | 18.4171 | 24.0000 | 0.7674 |  |  |
| Total | 30.1983 | 26.0000 |  |  |  |
| Pair-wise comparison with Bonferroni correction |  |  |  |  |  |
| Group 1 | Group 2 |  | Group 1 - Group 2 (95% CI) |  |  |
| NHP-A/Target Plan 1 | NHP-A/Target Plan 2 |  | -0.0643 (-1.1271, 0.9985) |  |  |
| NHP-A/Target Plan 1 | NHP-B/Target Plan 1 |  | -1.4323 (-2.4951, -0.3695) |  |  |
| NHP-A/Target Plan 2 | NHP-B/Target Plan 1 |  | -1.3680 (-2.4308, -0.3052) |  |  |

**Table 3:** Comparing MDPE across 2 NHPs and 2 target plans. One-way ANOVA table and pairwise comparison table reported. Each of the 3 groups comprised of 9 values of MDPE = 3 per experiments 3 × repeat experiments. SS - sum of squares, df - degree of freedom, MS - mean squared errors, F-estimated F-statistic which is the ratio of the mean squared errors, *Prob > F* denotes the probability that the ratio of the mean squared errors exceeds the estimated F-statistic, CI - confidence interval

| Source | SS | df | MS | F | $Prob > F$ |
| --- | --- | --- | --- | --- | --- |
| Columns | 1.8600 | 2.0000 | 0.9300 | 1.1675 | 0.3282 |
| Error | 19.1183 | 24.0000 | 0.7966 |  |  |
| Total | 20.9783 | 26.0000 |  |  |  |

Table 3: Comparing MDPE across 2 NHPs and 2 target plans.
| Group 1 | Group 2 | Group 1 - Group 2 (95% CI) |
| --- | --- | --- |
| NHP-A/Target Plan 1 | NHP-A/Target Plan 2 | 0.3377 (-0.7452, 1.4205) |
| NHP-A/Target Plan 1 | NHP-B/Target Plan 1 | 0.6426 (-0.4402, 1.7255) |
| NHP-A/Target Plan 2 | NHP-B/Target Plan 1 | 0.3050 (-0.7779, 1.3878) |

**Table 4:**
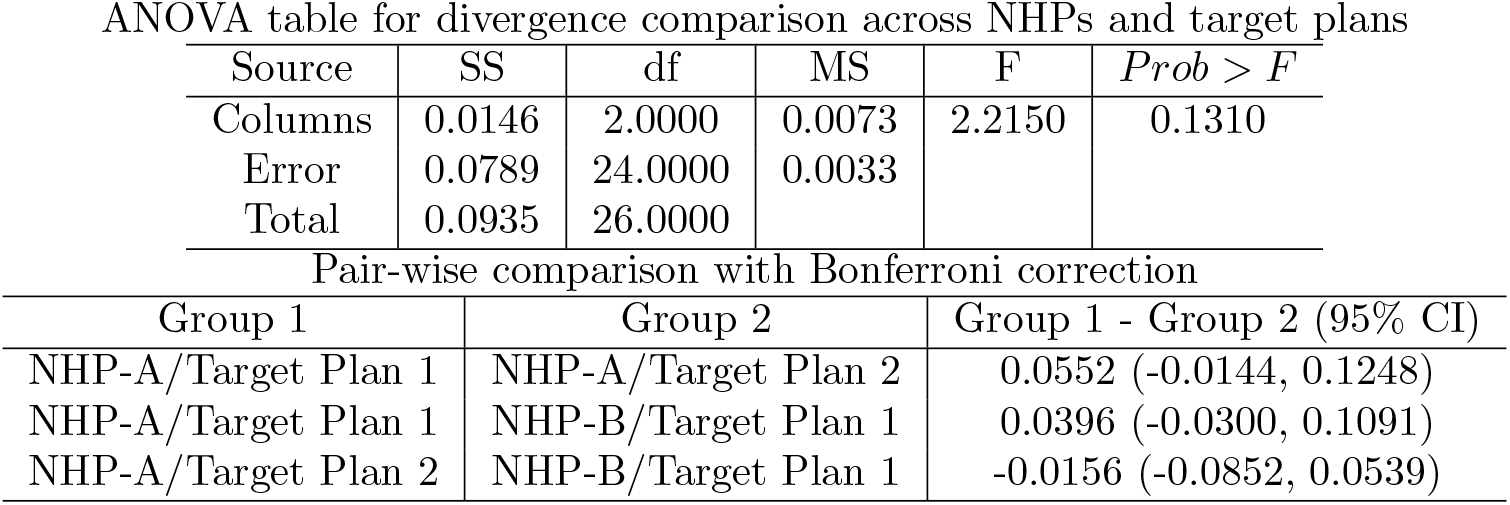
Comparing divergence across 2 NHPs and 2 target plans. One-way ANOVA table and pairwise comparison table reported. Each of the 3 groups comprised of 9 values of divergence = 3 per experiments × 3 repeat experiments. SS - sum of squares, df - degree of freedom, MS - mean squared errors, F-estimated F-statistic which is the ratio of the mean squared errors, *Prob > F* denotes the probability that the ratio of the mean squared errors exceeds the estimated F-statistic, CI - confidence interval

**Table 5:**
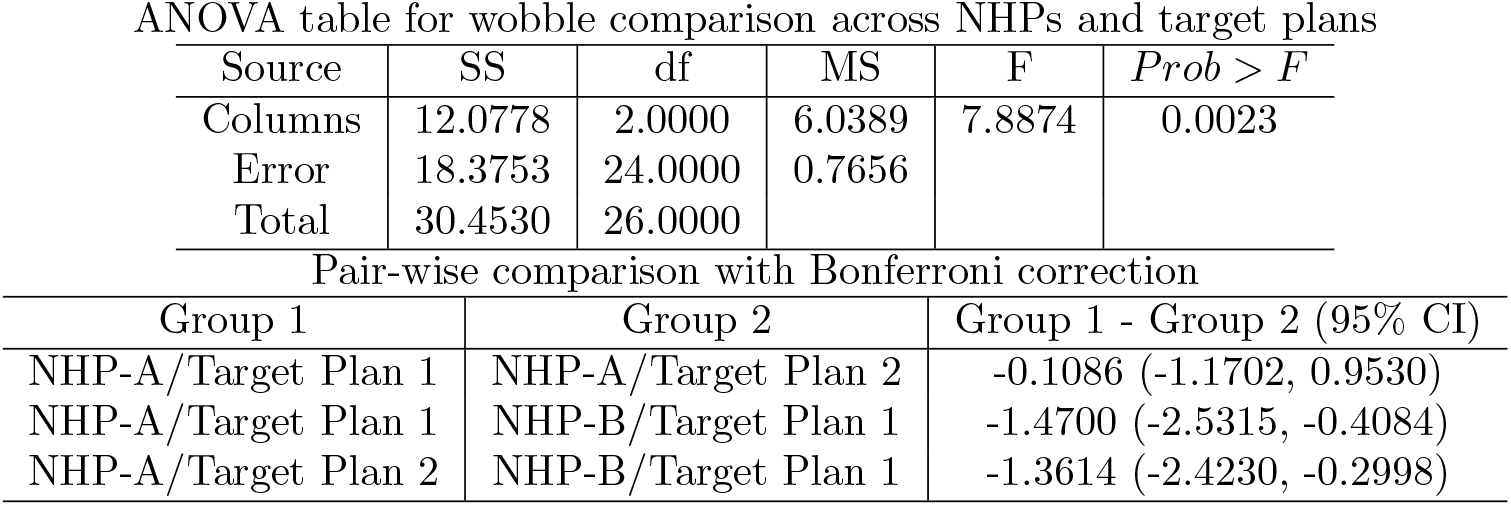
Comparing wobble across 2 NHPs and 2 target plans. One-way ANOVA table and pairwise comparison table reported. Each of the 3 groups comprised of 9 values of wobble = 3 per experiments 3 × repeat experiments. SS - sum of squares, df - degree of freedom, MS - mean squared errors, F-estimated F-statistic which is the ratio of the mean squared errors, *Prob > F* denotes the probability that the ratio of the mean squared errors exceeds the estimated F-statistic, CI - confidence interval

**Table 6:** Comparing average infusion rates across 2 NHPs and 2 target plans. One-way ANOVA table and pairwise comparison table reported. Each of the 3 groups comprised of 9 values of infusion rate averages = 3 per experiments × 3 repeat experiments. SS - sum of squares, df - degree of freedom, MS - mean squared errors, F-estimated F-statistic which is the ratio of the mean squared errors, *Prob > F* denotes the probability that the ratio of the mean squared errors exceeds the estimated F-statistic, CI - confidence interval

| Source | SS | df | MS | F | $Prob > F$ |
| --- | --- | --- | --- | --- | --- |
| Columns | 0.0090 | 2.0000 | 0.0045 | 2.1193 | 0.1420 |
| Error | 0.0512 | 24.0000 | 0.0021 |  |  |
| Total | 0.0602 | 26.0000 |  |  |  |
| Pair-wise comparison with Bonferroni correction |  |  |  |  |  |
| Group 1 | Group 2 |  | Group 1 - Group 2 (95% CI) |  |  |
| NHP-A/Target Plan 1 | NHP-A/Target Plan 2 |  | -0.0430 (-0.0991, 0.0130) |  |  |
| NHP-A/Target Plan 1 | NHP-B/Target Plan 1 |  | -0.0107 (-0.0667, 0.0453) |  |  |
| NHP-A/Target Plan 2 | NHP-B/Target Plan 1 |  | 0.0324 (-0.0237, 0.0884) |  |  |

**Table 7:**
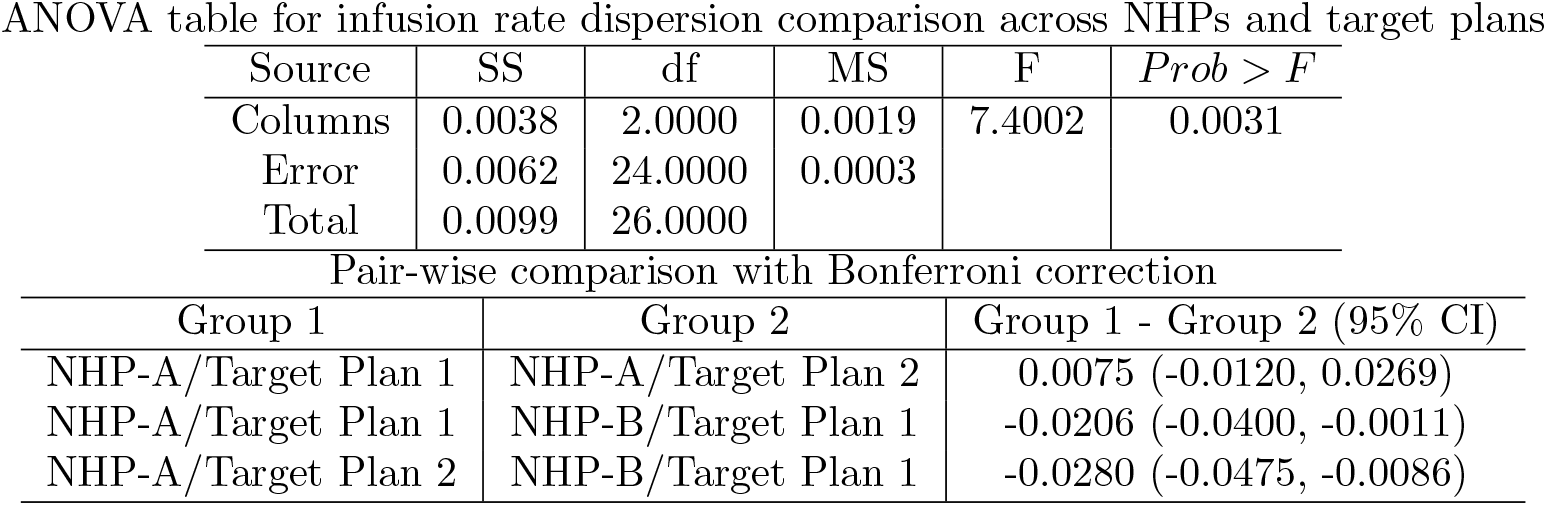
Comparing dispersion in infusion rates across 2 NHPs and 2 target plans. One-way ANOVA table and pairwise comparison table reported. Each of the 3 groups comprised of 9 values of dispersion in infusion rates = 3 per experiments× 3 repeat experiments. SS - sum of squares, df - degree of freedom, MS - mean squared errors, F-estimated F-statistic which is the ratio of the mean squared errors, *Prob > F* denotes the probability that the ratio of the mean squared errors exceeds the estimated F-statistic, CI - confidence interval

We can further analyze how control effort varied as the target MOU level switched between a low and a high target MOU level within a session. During an experiment with target plan 1 in NHP-A and NHP-B, as expected the average infusion rates does increase as the MOU target level is set to a lower value. This is indicated by the fact that the 95% CI, (-0.0945, -0.0392), of the mean pair-wise difference in *u*_*avg*_ between L1 segments (high MOU value) and L2 segments (with low MOU value) in Expts. 1-3 and 7-9 (Table 1) does not include zero. In the same experiment with target plan 1 when the target MOU is changed back to the same higher value as the first segment, as expected, the average infusion rate is lowered. This is indicated by the 95% CI, (0.0138, 0.0650), of the mean pair-wise difference in *u*_*avg*_ between L2 segments (high MOU value) and L3 segments (with low MOU value) does not include 0. However, interestingly, *u*_*avg*_ values from segments L3 are higher than L1 although both of segments have identical user-prescribed target MOU values. This is indicated by the 95% CI, (0.0076 0.0791), of the mean pair-wise difference between L3 segments and L1 segments across all the 9 experiments (Expt. 1-9 in Table 1) not including zero. This suggests that the equilibrium point (per a PK-PD model) corresponding to the same target MOU has changed, thus further suggesting that the PK-PD model itself has changed during the course of the propofol infusion session. Nevertheless, the CLAD system’s ability to maintain the MOU at similar levels of MDAPE, MDPE, divergence and wobble between L1 and L3 segments speaks to the robustness of the CLAD system in a long session with intra-session variability due to changes in the NHP’s neurophysiological response to propofol.

## 3 Discussion

Our work offers innovations on multiple fronts. For the first time, to the best of our knowledge, our work establishes a non-human primate (NHP) model to develop and test CLAD systems for maintenance of unconsciousness. The NHP model uses rhesus macaques and this provides (i). pre-clinical research and development paradigm (for CLAD) in an animal system whose brain and central nervous system are closely homologous to that of humans; and (ii). the animal model further allows repeated, systematic testing of the control markers and feedback control algorithms. Thus our work addresses a pre-clinical consideration recommended by FDA for closed-loop physiological system research [18]. The CLAD design and testing framework developed here can potentially be generalized to other anesthetic drugs. This opens up the scope for pre-clinical CLAD studies with other anesthetic drugs as well as combination of anesthetic drugs [36, 14, 37].

The experimental setup to record simultanoues LFP and spiking activity from an NHP’s cortex under propofol infusion enabled us to identify the normalized 20-30 Hz LFP power as the MOU to control by CLAD. This was based on our experimental finding that temporal changes in MOU were strongly correlated to changes in neuronal spiking activity (a neurophysiological correlate of unconsciousness during propofol-maintained anesthesia in NHPs [24, 23]). As a complementary side note, the bi-phasic trend in the MOU (initial rise followed by monotonic decline under a constant infusion rate) is also seen in the dynamics in beta frequency band (13-24 Hz) power of the scalp EEG in human studies with propofol anesthesia [7]. Further-more, the eletrophysiological recording setup used here is typical of NHP-based experimental neuroscience research. Also, the signal processing step to calculate MOU from the LFP signal using multitaper spectral analysis is also straightforward and can be executed with open-source algorithms [38]. Therefore, our choice of experimental setup and the MOU definition opens up opportunities to innovate further with neuroscience-informed CLAD research, and without facing the disadvantages due to lack of transparency in proprietary EEG monitoring algorithms that are currently used in clinical practice to monitor unconsciousness under anesthesia [39]. A limitation in our study is the lack of EEG recordings simultaneous with LFP and spiking activity recordings. A systematic scaling of our LFP-based CLAD system developed for NHPs to an EEG-based CLAD system in NHPs can be a natural next step that can support future EEG-based CLAD studies in humans. Another limitation of our setup was lack of behavioral data beyond eye-closed and eyes-open events. This was because the NHPs used here were subjects who were retired from previous non-CLAD experiments that were not designed to track changes in unconsciousness levels. This was intentional and motivated by safety consideration for the NHPs as this was the first time CLAD experiments were being performed in non-intubated NHPs. In future NHP-based CLAD research, it would be ideal to have continuous behavioral recordings under anesthesia (similar to prior NHP [23] and human [7] studies) from NHPs trained for unconsciousness research. This will allow for a more refined description of the level of unconsciousness and subsequently can improve the MOU definitions for CLAD purpose.

Our two-compartment PK and sigmoidal Emax PD models, estimated from concurrent neuroscience-informed MOU vs. infusion rate data, provides a pharmacologically-principled and accurate description of the MOU dynamics due to propofol infusion in a rhesus macaque. These NHP-specific PK-PD models were used for both model-informed feedback control design steps as well as for simulating MOU observations during *in silico* testing of candidate CLAD designs. Both the NHP CLAD design and testing framework, as well as the related findings, can inform future clinical studies of propofol CLAD with EEG feedback (see supplementary Sec B for a short discussion). A plausible and potentially feasible approach to extend the CLAD design reported here to a future human application, would be to replace the PK-PD model for NHPs with appropriate PK-PD models for humans, say e.g. the Schnider et. al.’s PK-PD model for propofol [40, 28]. The subsequent steps for the feedback control design and testing can remain the same, albeit with adjustments to incorporate the higher dimensional PK state and difference in parameter values. Another limitation in our study was that for the NHP-specific CLAD design the linearized system was based on a PK-PD model estimated from a single anesthesia experiment of the same NHP. It is important to note, that despite this limitation, our simulation and actual CLAD experiments demonstrate that our NHP-specific CLAD system can deliver stable performance with acceptable performance measures when there is model misspecification. A next step of this research could be to estimate NHP-specific PK-PD models from multiple experiments with the same NHP.

Our choice of the optimal robust control framework is a new CLAD innovation that combines the strengths of two well-validated principled feedback control strategies that has precedence in CLAD research - optimal control in rodent studies [16], and two-degree of freedom robust proportional integral control in human studies [31]. We efficiently constructed our CLAD algorithm by tailoring for CLAD the feedback control strategy proposed by Hagiwara et. al. [20, 21, 22] which allows flexibility in trading off between competing properties of good target tracking and good disturbance rejection under practical conditions of model misspecification and unmodelled disturbances. Prior to experimental deployment, we verified that the NHP-specific CLAD designs demonstrated acceptable performance in simulation studies. In these simulation studies we incorporated CLAD-specific practical conditions such as nonlinear PK-PD model, model misspecification, boundedness of infusion rate, and random noise in observed MOU. This systematic approach was aligned with the recommendations by FDA for such pre-clinical studies [18]. The modular nature of the analytic and experimental framework can allow for future studies where different feedback control strategies are tested in NHPs for the same MOU definitions and identical experimental conditions. Since our data suggests that the PK-PD model may be changing during an infusion session, an adaptive control paradigm (similar to Yang et. al. [41]) might be relevant in future extensions of the NHP-based CLAD system.

The experimental findings (Fig. 5, Table 1) establish the feasibility of using our CLAD system to stably, accurately and robustly control level of unconsciousness maintained by propofol at precise levels in rhesus macaques. Across, 1125 minutes (125 min/experiments × 9 experiments) the CLAD system is able to stably achieve and maintain target MOU values at acceptable levels across multiple sessions in a given NHP, across 2 NHPs and across 2 different target plans. The performance measures are deemed acceptable since they are comparable or lower than those reported in literature (Table 8 provides a few representative numbers for performance measures reported in CLAD literature). Our findings therefore advances CLAD research closer towards future clinical studies in the USA. Another scope for further research would be to test the CLAD systems in NHPs who are undergoing elective surgeries. This prospective surgical scenario can allow for testing the CLAD under more realistic conditions of noxious stimuli and with other drugs on board. In such prospective surgical scenario, since the NHPs will most likely be intubated and be under veterinary supervision, relaxation of the upper limit on the infusion rate might be feasible and thereby potentially improving on CLAD performance and without compromising on safety.

**Table 8:** Performance measures from a few previous studies on propofol-based CLAD. Numbers reported are those quoted in the corresponding publication. The representation *x* ± *a* indicates mean ± standard deviation, whereas the representation *x*(*a, b*) indicates the median (25th percentile, 75th percentile) where the central tendency and dispersion parameters are calculated across multiple patients or animal subjects. All central tendency measures that are reported without any associated uncertainty measure are reported as medians.

| Publication | Country | Model | MDAPE (%) | MDPE (%) | Divergence (%/min) | Wobble (%) |
| --- | --- | --- | --- | --- | --- | --- |
| Struys et al. [35] | Belgium | Patients | $7.7 \pm 2.49$ | $-6.6 \pm 2.63$ | $0.024 \pm 0.029$ | $5.90 \pm 2.33$ |
| Absalom et al. [10] | United Kingdom | Patients | 8 | 2.2 |  | 7.3 |
| Liu et al. [13] | France | Patients | $9.94 \pm 3.4$ | $-3.32 \pm 5.37$ | $8.10 \pm 2.46$ | |
| West et al. [11] | Canada | Patients | 8.4 (5.1, 12.1) | -4.6 (-10.5, 2.2) |  | 5.2 (4.2, 7.1) |
| Puri et al. [15] | India | Patients | 10 (10, 12) |  |  | 9 (8, 10) |
| Shanechi et al. [16] | USA | Rodents | 3.61 | -1.44 |  |  |

In summary, our work establishes an NHP-model for CLAD. This CLAD system estimates an animal’s level of unconsciousness in real-time based on an LFP-based MOU and then uses the error between the estimated MOU and a user-prescribed target value to titrate the propofol dose so that the observed MOU tracks the user-prescribed target MOU value. The automatic LFP feedback-based titration is achieved using a feedback control framework that is applied for the first time in a CLAD application. The feedback control framework is tuned for each NHP using NHP-specific parsimonious PK-PD model. Our experimental studies demonstrate that the CLAD system developed here provided stable and accurate performance and was robust to intra-experiment, inter-repeat (for same subject and target plan), inter-subject and inter-target plan variabilities. Our systematic study in NHPs advances CLAD research and further reinforces the feasibility of CLAD to safely and accurately maintain unconsciousness in patients for prolonged duration. Thus, this research provides a critical foundation needed to develop and deploy CLAD systems for use with patients requiring anesthesia care in the operating room or in the intensive care unit.

## 4 Materials and Methods

### 4.1 Experimental paradigm and recording setup

Two rhesus macaques (macaca mulatta) were chronically implanted with multi-electrode arrays (MultiPort Utah Array, Blackrock Microsystems, Salt Lake City, UT) placed in the prefrontal cortex. In subject NHP-A (male), we simultaneous recorded from arrays in supplemental eye field (SEF) and dorsolateral PFC (dlPFC). In subject NHP-B (male), we simultaneously recorded from arrays in dlPFC and ventrolateral PFC (vlPFC). From each subject, both raw signal at 30 KHz and a filtered 1 KHz signal low-passed below 250 Hz were recorded. Monkeys were head-fixed in a sitting posture via an implanted titanium head-post and placed in a noise-isolation chamber. In each subject, prior to an experimental session, vascular access was established through an ear-vein cannula for anesthesia administration (cannulation performed by NHP anesthesia expert (MM)). During an anesthesia session, the propofol (commercially available in 10 mg/ml concentration) was infused via programmable a syringe pump (PHDUltra, Harvard Apparatus, Holliston, MA). The flow rates were prescribed using a custom Matlab script communicating with the pump through a serial port using functionalities from Matlab’s Instrument Control Toolbox (running on the same computer that is also running the Blackrock Central data acquisition software suite). During the CLAD sessions, the 1 KHz signal was made accessible within the Matlab workspace by leveraging the Blackrock’s *cbmex* interface. LFP recordings from a pre-selected single electrode were used for CLAD algorithm design and experimental testing. The single electrode was selected based on the observation by an NHP electrophysiology expert of multi-unit activity characterized by distinct spike waveforms prior to experimental testing of the CLAD algorithm and then kept fixed for all CLAD sessions in each subject. Infrared monitoring tracked facial movements, and pupil size (Eyelink 1000Plus, SR-Research, Ontario, CA) throughout the course of the experiments. Physiological monitoring of heart rate and oxygen saturation was performed by dedicated NHP anesthesia experts (ASW, MM) throughout the period of the recording to ensure safety of the animal (using Model 7500, Nonin Medical, Inc., Plymouth, MN). Occasionally, when oxygen saturation approached 90%, breathing support was provided manually using an ambu bag. Throughout each experiment session, we ensured that the oxygen saturation stayed steadily above 90% and subject was breathing on its own without intubation.

### 4.2 Spectral analysis of single-channel LFP from a given recording session

In both offline and online (realtime) spectral analysis of 1 KHz single-channel LFPs, we used non-overlapping, consecutive data windows of Δ = 20 seconds = 0.3 min. We estimated the instantanoues power spectra using the multitaper spectral analysis (Chronux Toolbox [38]) with the time-halfbandwidth product = 4 and number of tapers = 3. To calculate MOU for a given Δ interval, first the LFP power (in dB) in 20 – 30 Hz range was calculated from the corresponding multitaper spectrum. Then, the MOU estimate was determined by dividing this LFP power in 20-30 Hz band by a normalization factor. The normalization factor here was the median of power from the same band determined from a 5 minute period of data immediately prior to start of intravenous propofol infusion. In the offline setting, a smoothed estimate was obtained by fitting a cubic smoothing spline to the entire MOU sequence using Matlab function *csaps*() with its smoothing parameter set as 1*/*(1 + Δ^3^*/*0.06). This smoothed estimate was used for both visualizing the LFP marker trend as well as estimating the session-specific PK-PD models.

### 4.3 Spike rate calculation from a given recording session

The spike rates were calculated in an offline setting by post-processing the 30 KHz data from multiple channels in the same microelectrode array from which we selected the single electrode for MOU calculation (Sec 4.2). First, we band-passed the signal from each electrode between 250 Hz to 5000 Hz using a second-order Butterworth filter. Then we determined standard deviation of one-minute segment of the signal from the middle of the anesthesia session. We used a threshold-based approach to detect spiking where we identified all voltage activity whose magnitude exceeded 4.5× the estimated standard deviation. Then we explicitly imposed a 1.5 millisecond refractory period to identify a binary time-series of inferred spiking activity (such that 1 would indicate a single spike event, and 0 otherwise). From such binary time-series from significantly spiking electrodes (whose total spike counts across the entire session exceeded 90-th percentile from all channels in the relevant microelectrode array), we calculate the ensemble spike rate in the *k*-th Δ interval by summing up all the 1’s across all the channels from that interval and converting them to dB scale. Finally, to obtain the baseline normalized spike rate we divide spike rate sequence by the median value from the 5 min interval immediately prior to start of propofol infusion. For visualization purpose only, and similar to the approach used for the LFP MOU data (Sec. 4.2), we estimate a smoothed spike rate sequence using Matlab function *csaps*() with its smoothing parameter set as 1*/*(1 + Δ^3^*/*0.06).

### 4.4 PK-PD model and related parameter estimation from a given anesthesia recording session

Mathematically, the PK component of the PK-PD model described above is given by the following continuous-time state-space equation (in modal form),

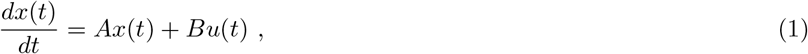

where, *x*(*t*) = [*x*_1_(*t*), *x*_2_(*t*)]^*T*^ with the superscript [·]^*T*^ denoting a matrix transpose operation, *A* is a diagonal matrix with *A*_1,1_ = *λ*_1_ and *A*_2,2_ = *λ*_2_ with *λ*_2_ *< λ*_1_ *<* 0, and *B* = [1, 1]^*T*^. The amount of drug in the effect compartment is given by,

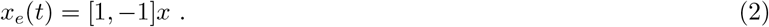

The sigmoidal function describing the decreasing trend in the MOU, *y*(*t*), with increasing *x*_*e*_(*t*) during the period of unconsciousness is posited to be,

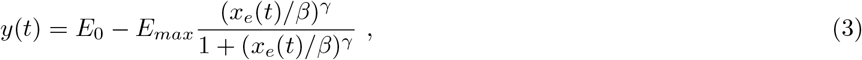

where, *E*_0_, *E*_*max*_, *β* and *γ* are positive scalar-valued parameters. To be precise, for a two-compartment mamillary model (Fig. 3(A), top panel) *x*_*e*_ ∝ [1, −1]*x*, but we set it as an equality since the proportionality factor cannot be distinguished from *β* when we seek to estimate all the PK-PD model parameters directly from *u*(*t*) and *y*(*t*) data (see [42] and [29, Sec. C] for details). Therefore, the parameter set Θ_*P*_ = {*E*_0_, *E*_*max*_, *γ, β, λ*_1_, *λ*_2_} characterizes a PK-PD model.

To estimate Θ_*P*_ from a given recording session of duration *K*Δ, we fit the PK-PD model to the available information on 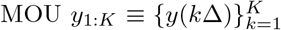 at every Δ time interval and infusion rate history upto the same duration, {*u*(*t*) : 0 *< t < K*Δ}. To minimize the effect of the outliers in the optimization problem we fit the model to the smoothed estimate of LFP marker 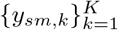derived from *y*_1:*K*_ (see Sec. 4.2 for details on the smoothing operation). First we set *E*_0_ to be the maximum value that the MOU attains before starting its descent during a monotonically non-decreasing infusion rate of propofol. To estimate the other PK-PD parameters, we minimize the following objective function,

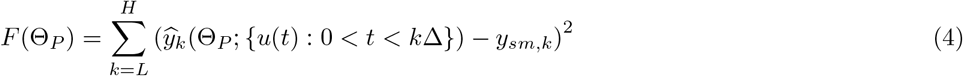

where, *ŷ*_*k*_ denotes the model predictions at time-point *k*Δ calculated by substituting Θ_*P*_ and infusion rate history {*u*(*t*) : 0 *< t < k*Δ} in Eqs. (1), (2) and (3). The time-point *k* = *L* is taken to be the EC time-point. The time interval [*L*Δ, *H*Δ] corresponded to the period of unconsciousness. By our choice of *L* and *H*, note that our PK-PD model will be valid in our anesthetic regime of interest where the LFP marker decreases with increase in *x*_*e*_. Starting with an initial guess, we estimate the parameters using an alternating optimization approach where we alternate between the two minimization problems to estimate the PD and PK parameters separately (similar in principle to our earlier work[29]).

The initial guesses for the PD parameters are given by 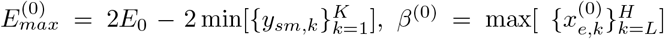, *γ*^(0)^ = 1, where superscript ((·)^(0)^) denotes the 0-th iteration of the alternating optimization. The initial guesses for the PK parameters, 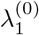 and 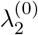, are set as the maximum and minimum eigenvalues of the transition matrix characterizing the PK of the two-compartment model that is derived from the 3-compartment model reported earlier in the following manner [43]. In the absence of a propofol PK model for rhesus macaques, we consider the three compartment linear PK model determined for Japanese macaques [43]. We derived a reduced-order (a two-compartment mamillary model [42]) that has an impulse response function for the effect compartment most similar to the impulse response function for the central compartment of the three-compartment mamillary PK model ([43]).

In the *i*-th iteration of the iterative alternating optimization approach, we first solve the following constrained optimization problem using Matlab’s *fmincon()* function,

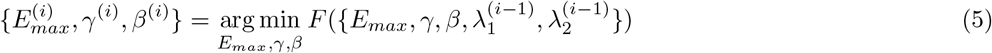

such that 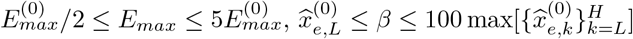, where,

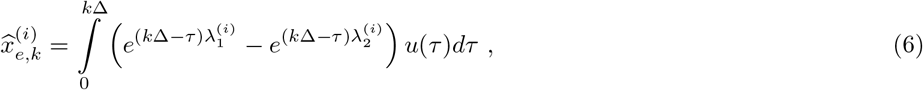

and 0.1 ≤ *γ* ≤ 20. After solving the optimization problem in Eq. 5, we solve another optimization problem

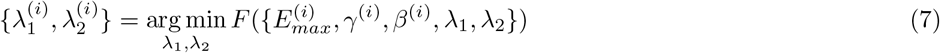

such that −10000 *< λ*_1_ *< λ*_2_ *<* 0. We alternate between the two optimization problems, Eqs. (5) and (7), for a prescribed maximum number of iterations (50 iterations in our implementations). Finally, we report the inaccuracy between the estimated MOU *ŷ* and the given data *y* using a relative error metric calculated as 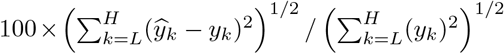.

### 4.5 Controller design parameters for a given subject

In choosing our control strategy we considered existing strategies reported in literature that can enable the output of the system being controlled (the MOU response in the current case) to track a user-prescribed target value (target MOU). Those strategies were favored that were similar to ones that have precedence in the CLAD literature, such as the optimal control framework (Linear Quadratic Regulator (LQR) strategy with estimated state feedback) for CLAD as in our prior rodent studies[16, 17] and the PID control framework for CLAD as used in human subjects outside the USA [31, 32, 15]. In a separate simulation-based study we verified that a hybrid approach, with the principled optimization-based design framework of the former and the stable output tracking property under model uncertainty of the latter, could stably control the pharmacodynamic response of propofol PKPD models of human subjects[33]. This inspired us to choose the optimal robust servo-control scheme that updates the infusion rate by combining two components: an LQR strategy and an integral compensation strategy that monitors accumulation of the tracking error (*MOU*_*target*_|*MOU*_*actual*_)[20, 21, 22]. The LQR component uses an observer to compute the propofol effect-site level[34, 22]. Since this LQR, when coupled to an estimator, can be sensitive to model mis-specification, we included an integral compensation that can ensure stable MOU tracking even when the PK-PD model is mis-specified. Hence, the integral compensation helps ensure a degree of robustness in our control strategy. In addition to the aforementioned benefits, the optimal robust servo-control framework provides a principled optimization-based framework to design a two-degree of controller for a linear system with step targets and step disturbances. The “two-degree of freedom” feature of the controller stems from the fact that the response to step responses and to step disturbances can be tuned separately without affecting the performance of the either. For the optimal robust servo-control this is done by minimizing two distinct quadratic criteria6. The solution of the first leads to the LQR feedback gains and the solution to the latter characterizes the integral action and observer gains. The control design is characterized by the set of user-prescribed parameters that goes in the definitions of the quadratic criteria.

Choosing the optimal robust servo control strategy is just the first step towards our CLAD controller design. Tuning the parameters of the controller such that the CLAD has desired properties was the next step. Since the optimal robust servo control strategy was developed for linear systems, we linearized a PK-PD model (estimated from an NHP) about a steady state infusion rate of *u*_*ss*_ = 0.285 mg/kg/min, a value that we determined empirically to be a safe dose for maintaining unconsciousness during a prolonged experimental session for the NHP. The steady-state values for the PK state, *x*_*ss*_ and for the MOU, *y*_*ss*_ were calculated as follows. According to the PK model (Eq. (1) and (2)) is given by

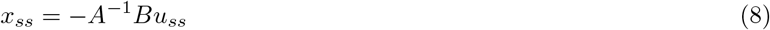

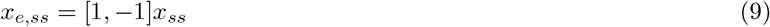

Then steady state response *y*_*ss*_ is calculated by substituting *x*_*e*_ with *x*_*e,ss*_ in Eq. (5). Assuming that the CLAD operation will be primarily in the neighborhood of this steady state, the next step was to derive an linear system approximation that captured the deviation of the state and MOU about the steady state.

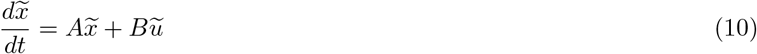

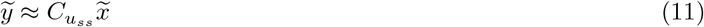

where, 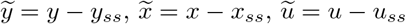, where 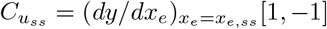.

From this step onwards, the feedback control problem was rephrased as that of output tracking problem (where 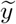 should track a constant target value, 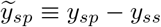) of a linear time-invariant system with additive disturbance terms, *d*_*x*_ and *d*_*y*_.

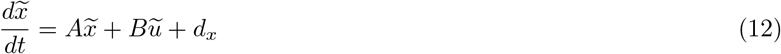

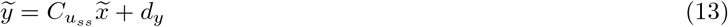

This pose of the feedback control problem with system dynamics given by Eq. (12) and (11) was amenable to controller design per Hagiwara et al.’s two-degree of freedom optimal robust control framework [20, 21, 22]. The optimal feedback control, *u*_*c*_, is calculated as

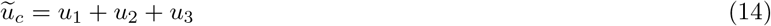

where, *u*_1_ is an output of a LQR control block, *u*_2_ is a feed-forward value that would account for non-zero setpoint, 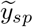, and *u*_3_ corresponds to the control contribution due to integral compensation. The LQR control is given by,

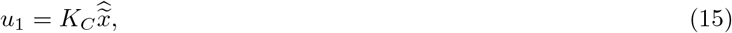

where, *K*_*C*_ denotes the LQR gain and 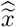 denotes the estimated deviation of the state from its nominal value *x*_*ss*_. Here, the LQR gain, *K*_*C*_, can be calculated as,

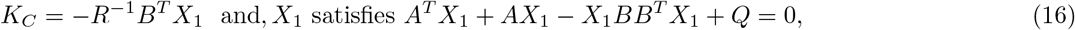

where, *Q* and *R* are user-prescribed parameters to enforce a design choice specifying contributions of deviations in state trajectory and deviation in control trajectory, respectively, to a quadratic criterion that is minimized [21, 44, 16]. We further impose structure on the parameters *Q* and *R* as follows,

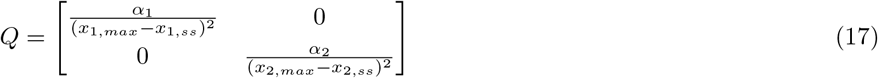

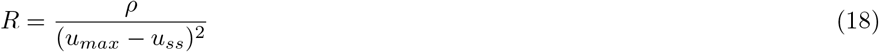

where, *x*_1,*max*_ and *x*_2,*max*_ are steady state values corresponding to a user-prescribed *u*_*max*_ (=4 mg/kg/min in this work). Therefore, LQR design is characterized by *α*_1_, *α*_2_, and *ρ*. The dynamics of the estimate 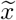 is governed by the following differential equation,

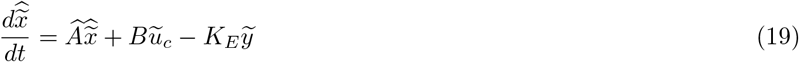

where, 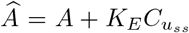 [34]. A stable estimator design can be specified by choosing the estimator gain, *K*_*E*_, such that *Â* has negative eigenvalues. Following Hagiwara [22], in this work, *K*_*E*_, is prescribed as Kalman Filter-like gain, per,

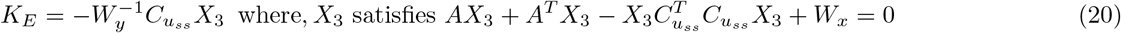

Again, *W*_*x*_ and *W*_*y*_ are design parameters that can be regarded as terms that capture the variability in the process dynamics and the measurements, respectively (similar in principle to covariance matrices of random noise terms in the process and observation equation of a Kalman Filter[45]). Again following Hagiwara et al. [22], we impose the following structure on *W*_*x*_ and *W*_*y*_,

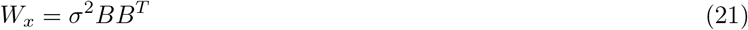

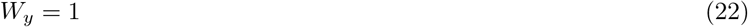

Therefore, specifying *σ* enforces a design choice for the estimator dynamics.

The feed-forward input *u*_2_ is given by,

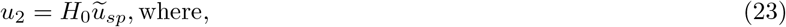

where, the feed-forward gain, *H*_0_ is given by,

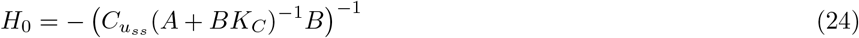

In the absence of disturbances and modeling errors, the term *u*_3_ = 0 in Eq. (14) is zero [21]. When disturbances and modeling errors are present, *u*_3_ acts as a correction term enabling output tracking via an integral compensation. The expression for *u*_3_ is given by,

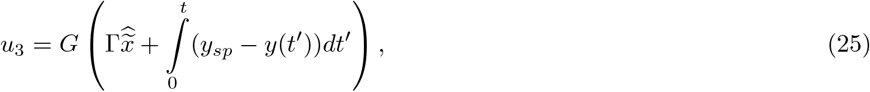

where, 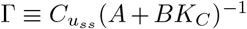. Note that *u*_3_ has contributions from both the estimator output and the integral of the steady state error. Following Hagiwara et al.[21], the gain *G* is given by,

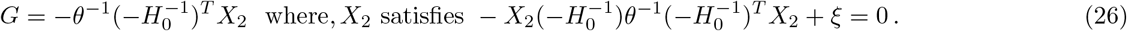

where, *θ* and *ξ* are user-prescribed positive-valued scalar design parameters that enforce design choices on disturbance rejection properties of the controller when disturbances and model misspecifications are present ^1^. Similar to the *Q* and *R*, the terms *ξ* and *θ* serve to enforce design choices for disturbance rejection properties in another quadratic criterion that is minimized to determine G per Eq. 26 [21].

The linear controller design for a given NHP is specified by the set of parameters, Θ_*C*_ = {*θ*_*P*_, *α*_1_, *α*_2_, *ρ, σ, θ, ξ*}. By leveraging the continuous-time framework of the optimal robust servo control strategy, one can determine the loop transfer function and consequently determine metrics that capture the robustness of the controller and convergence dynamics. This information can be determined by leveraging Matlab’s Control Systems Toolbox and using *margin()* and *stepinfo()* subroutines with the loop transfer function as an argument. The loop transfer function can be calculated as a product of two complex-valued functions *G*_*P*_ (*s*) and *G*_*Y*_ (*s*), where,

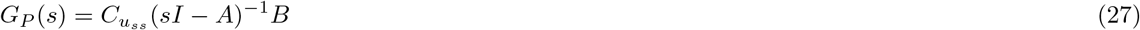

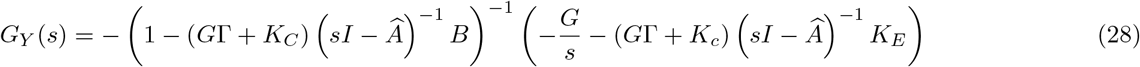

and *s* denotes a complex number.

In simulation studies, we tested the ability of this linear feedback control strategy to provide acceptable tracking performance when the dose-effect site relationship was governed by the NHP’s PK-PD model. To mimic actual experimental conditions in our simulations, we impose the lower and upper limits (*u*_*min*_ = 0 mg/kg/min and *u*_*max*_ = 0.4 mg/kg/min) on infusion rates on the suggested control *u*_*c*_ to determine the actual control signal *u* that is communicated to the infusion pump. Furthermore, we added Gaussian white noise (zero mean and standard deviation of 0.02 ; standard deviation value was empirically determined from the residual error between MOU and the estimated MOU using the best fitting PK-PD model) to the PK-PD model output, and enforced fixed rate of infusion between each discrete controller updates. For a given NHP’s PK-PD model, Θ_*P*_, we tuned the free parameters, {*α*_1_, *α*_2_, *ρ, σ, θ, ξ*} and tested each candidate controller design in *in silico* experiments that comprised a 125 closed-loop simulation with 3 step input changes at 0 min, 45 min, 85 min after a 30 min open-loop simulation under constant infusion rate of 0.285 mg/kg/min. Through this manual tuning exercise we selected that control design whose open-loop transfer function (−*G*_*P*_ *G*_*Y*_) had high phase margin (*>* 80 degree), high gain margin (*>* 60 dB) and low settling times for a unit step input (*<* 11 min), and the CLAD performance was acceptable. The performance criteria chosen was that of inaccuracy being less than 5% across the 125 min of closed-loop simulation. Briefly, a high gain margin for the control design indicates that the closed-loop system will still remain stable even when the true system has a multiplicative gain factor that was unaccounted during control design. The margin provides the extent to which this unaccounted gain factor can be tolerated. Similarly, a high phase margin would indicate that the closed-loop system would still remain stable even when the true system has delay (up to an extent specified by the margin) that was unaccounted during control design. Lower settling times for a unit step indicates faster convergence of MOU to target when a new target value is encountered. High gain and phase margins, and low settling times are desirable properties for a CLAD design.

### 4.6 Analysis of CLAD Performance

The controller performance was analyzed using the sequence of data {(*y*_*k*_, *u*_*k*_)} where *y*_*k*_ and *u*_*k*_ denoted the MOU value and the infusion rate at the time-point *t*_*k*_ (say, prescribed in minutes) where, *k* ∈ {1, …, *K*} with *k* = 1 and *k* = *K* denoting the beginning and the end of a temporal segment. Using this data and the definition of instantaneous performance error, *e*_*k*_ = 100(*y*_*k*_ − *y*_*sp*_)*/y*_*sp*_, several CLAD performance metrics that are typically used in literature can be calculated [46, 47, 31, 16]. These are metrics are: median absolute performance error (MDAPE), Median performance error (MDPE), Wobble (Wob), and Divergence (Div).

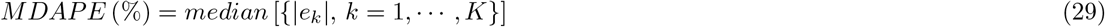

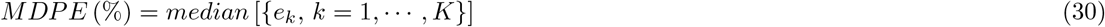

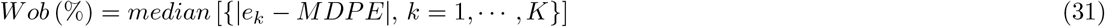

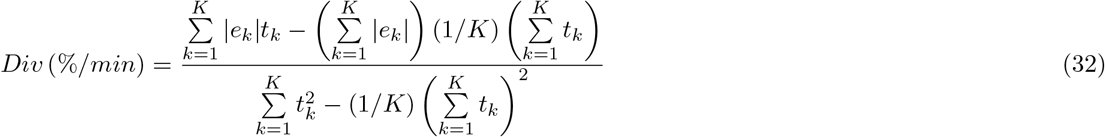

The *MDAPE* is a measure of the inaccuracy in CLAD performance for the entire period of observation. The *MDPE* is a measure of the bias in the MOU target tracking, with a positive (or, a negative) value indicating that the MOU lies mostly above (or, below) the target value. Wobble provides a measure of the intra-session time-related variability, with a high wobble value indicating a higher degree of oscillation of the controlled MOU about its target. Divergence characterizes any linear trend in the performance error over time, with a positive value suggesting that the MOU is gradually diverging away from the target, whereas a negative value suggesting a gradual converging of the MOU towards the target. Additionally, we also report the average (denoted by *u*_*avg*_) and dispersion (as measured by standard deviation and denoted by *u*_*std*_) of the infusion rates across the *K* data-points. The statistical comparison of the CLAD performance measures across multiple experiments or across target MOU levels (e.g. plotting of the boxplots, calculation of the ordered statistic, execution of the one-way ANOVA analyses and two-group comparisons) were performed using Matlab functions: *median(), prctile(), anova1(), multcompare()* and *ttest()*.

## 7 Acknowledgement

The authors acknowledge the help provided by Dr. Robert Marini, Dr. Alexis Garcia, Dr. Jefferson Roy, Dr. Scott Brincat, Dr. Jesus Ballesteros, Jordan DeFarias, and Anna Rock with animal surgery and animal care, and/or operational details towards experiments, data curation and data analysis. S.C. thanks Dr. Jingzhi An, Dr. John Abel, Dr. Gabriel Schamberg, Dr. Dimitrios Papageorgiou, Hugo Soulat, Devika Kishnan, Ksenia Nikolaeva, David Zhou, and John Tauber, for helpful discussions and for initial code snippets. S.C. further thanks Dr. Francisco Flores, Dr. Christa Van Dort, Dr. Patrick Purdon, Dr. Yumiko Ishizawa, and Dr. Michelle McCarthy for additional helpful discussions.

## 8 Author Contribution

**Conceptualization**: Sourish Chakravarty, Emery N. Brown, Earl K. Miller.

**Data curation**: Sourish Chakravarty, Ayan S. Waite.

**Formal analysis**: Sourish Chakravarty, Emery N. Brown.

**Funding acquisition**: Emery N. Brown, Earl K. Miller.

**Investigation**: Sourish Chakravarty, Jacob Donoghue, Ayan S. Waite, Meredith Mahnke.

**Methodology**: Sourish Chakravarty, Jacob Donoghue, Meredith Mahnke.

**Resources**: Emery N. Brown, Earl K. Miller.

**Software**: Sourish Chakravarty, Ayan S. Waite, Indie C. Garwood.

**Supervision**: Emery N. Brown, Earl K. Miller.

**Validation**: Sourish Chakravarty.

**Visualization**: Sourish Chakravarty.

**Writing – original draft**: Sourish Chakravarty, Emery N. Brown.

**Writing – review editing**: Sourish Chakravarty, Emery N. Brown, Earl K. Miller, Jacob Donoghue, Ayan S. Waite, Meredith Mahnke, Indie C. Garwood.

## A Supplementary information

**Figure 6:**
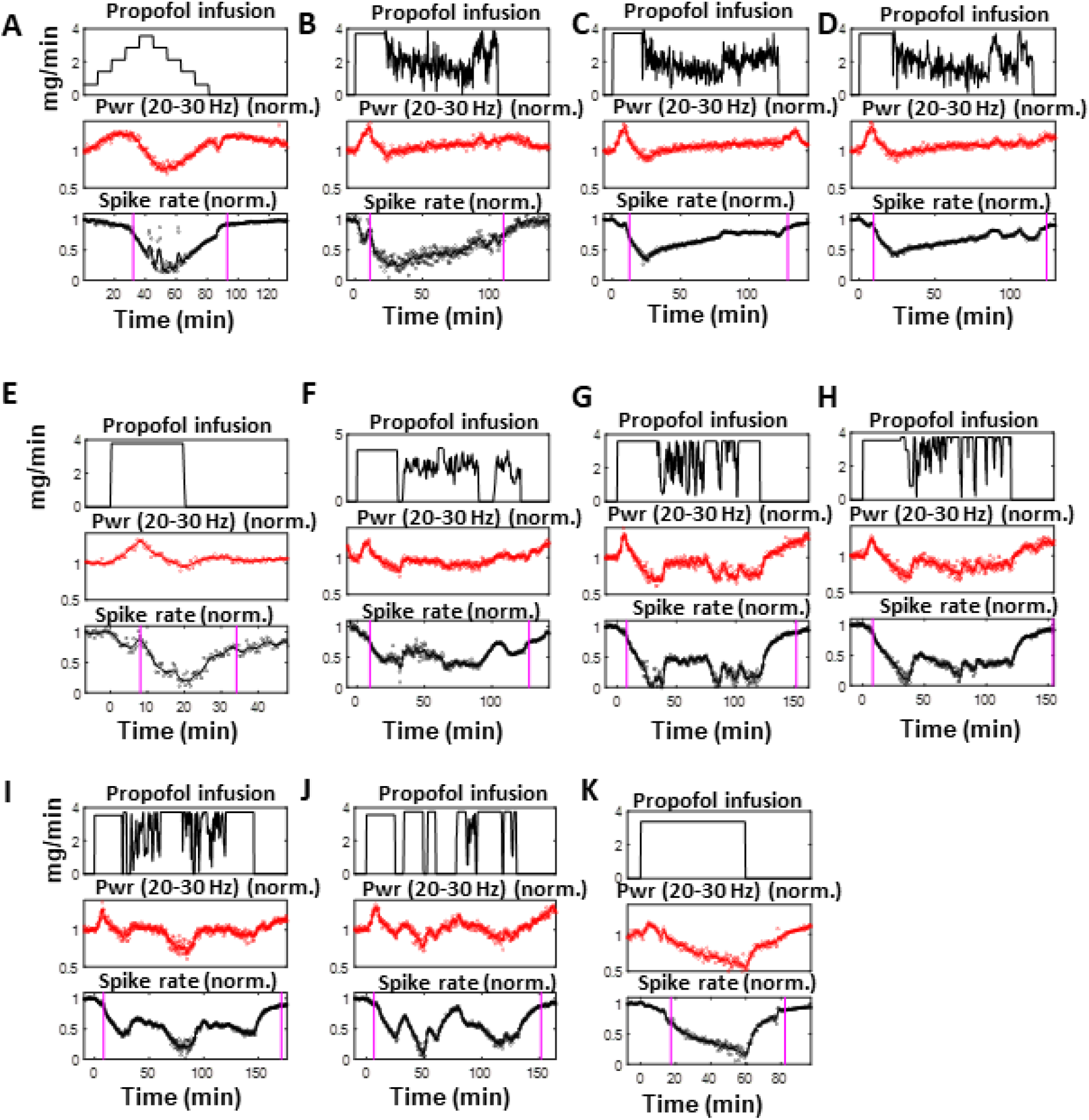
Neurophysiological response under across 10 anesthesia sessions in NHP-A (sub-figures A-K) and in NHP-B (sub-figure K). Each sub-figure (A through K) presents the infusion rate (top panel), MOU (middle panel), normalized spike rate (bottom panel) traces time from a distinct anesthesia session. The vertical magenta lines represents the EC and EO time-points in chronological order.

**Figure 7:**
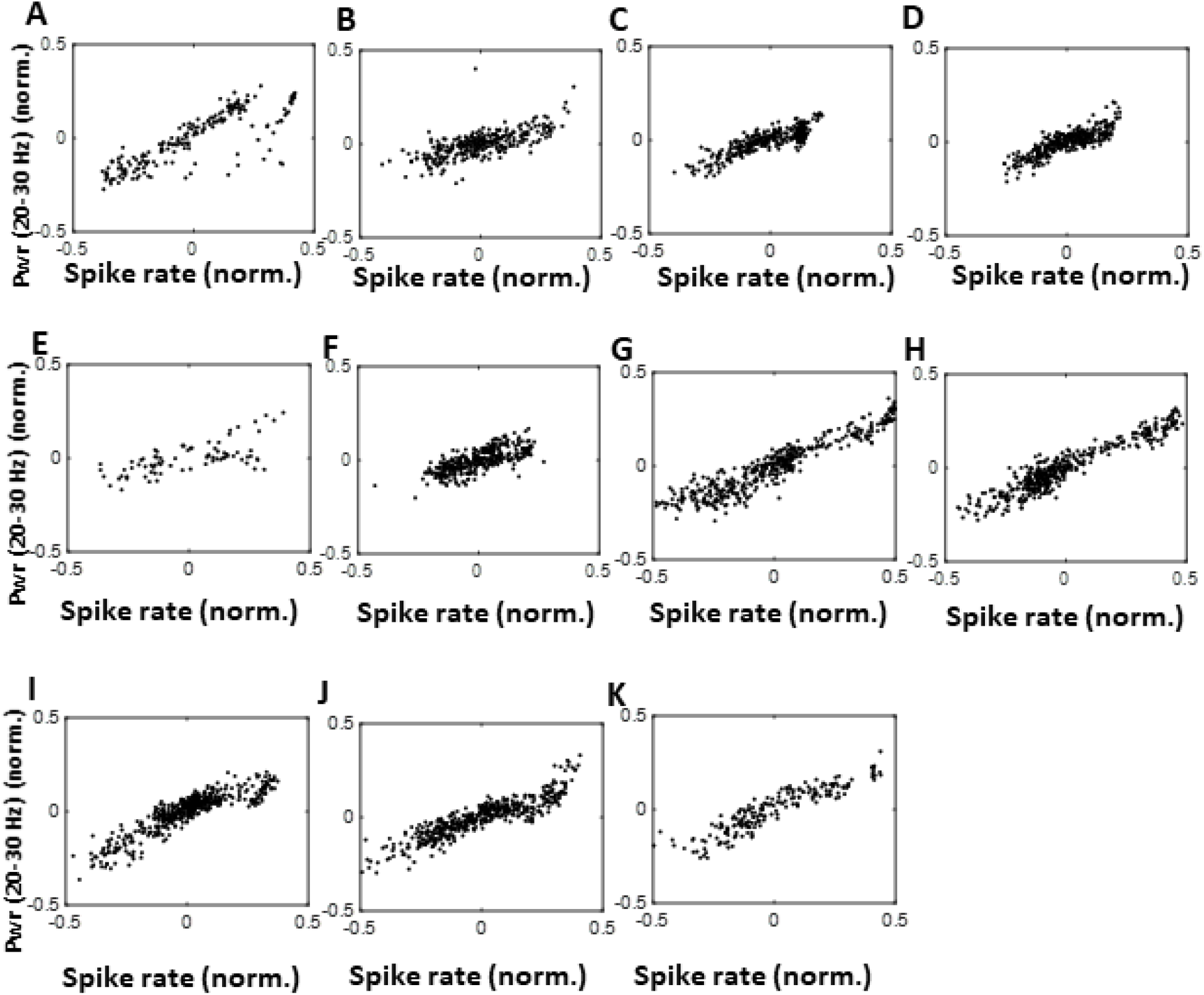
MOU vs normalized spike rate plots across multiple anesthesia sessions from the period of unconsciousness. The data is centered about its respective mean value. Sub-figure A through J are from 10 different anesthesia sessions from NHP-A. Sub-figure K is from the same anesthesia session in NHP-B that is reported in Fig. 2(A-C). Spearman’s rank correlation parameter estimate and their 95% confidence intervals for each session are given as follows. A: 0.8318 (0.8130, 0.8490), B: 0.7243 (0.7026, 0.7447), C: 0.8094 (0.7945, 0.8233), D: 0.8185 (0.8040, 0.8320), E: 0.6608 (0.6096, 0.7065), F: 0.7503 (0.7319, 0.7675), G: 0.9340 (0.9288, 0.9388), H: 0.9075 (0.9005, 0.9141), I: 0.8938 (0.8863, 0.9009), J: 0.9071 (0.9000, 0.9137), K: 0.9314 (0.9233, 0.9387)

**Figure 8:**
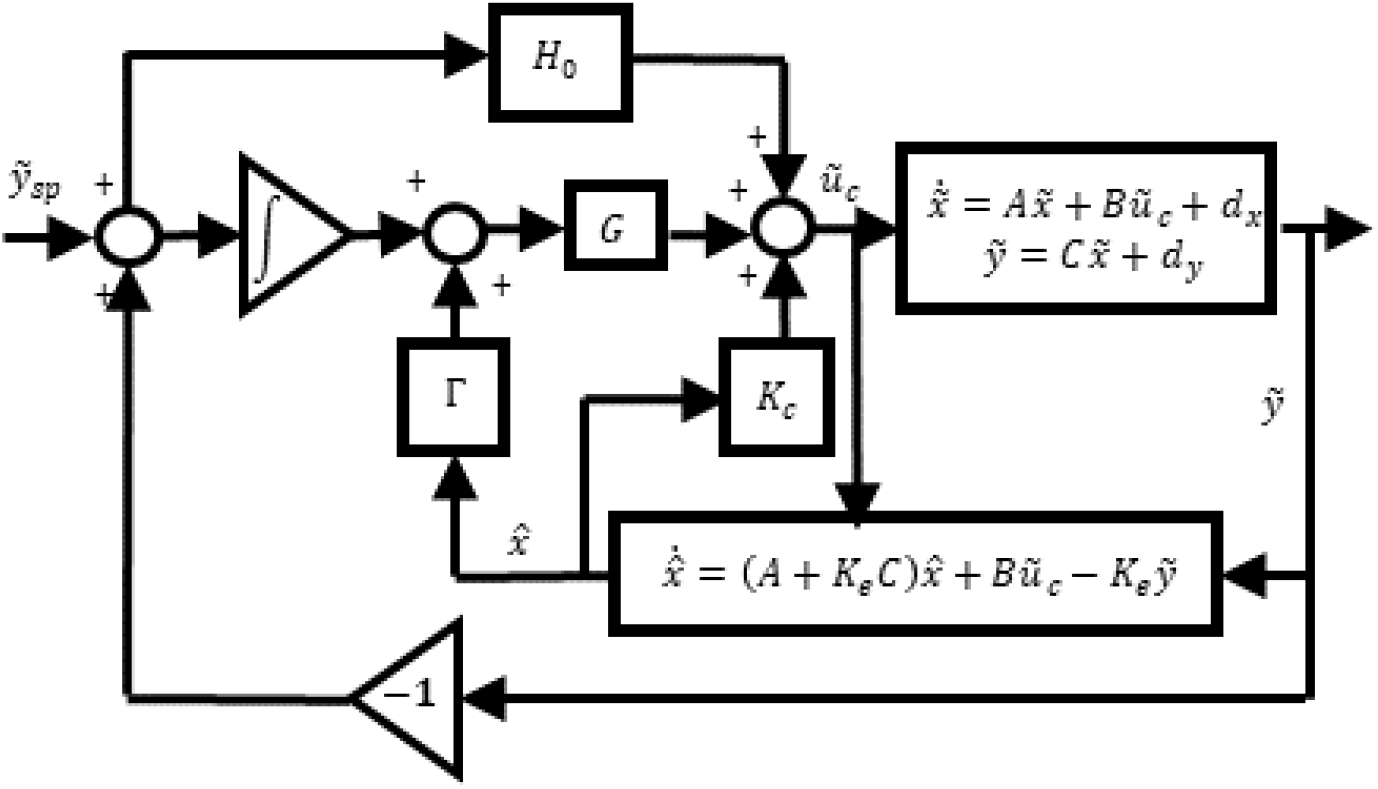
Detailed block diagram of the optimal robust servo-controller for the linearized system.

**Figure 9:**
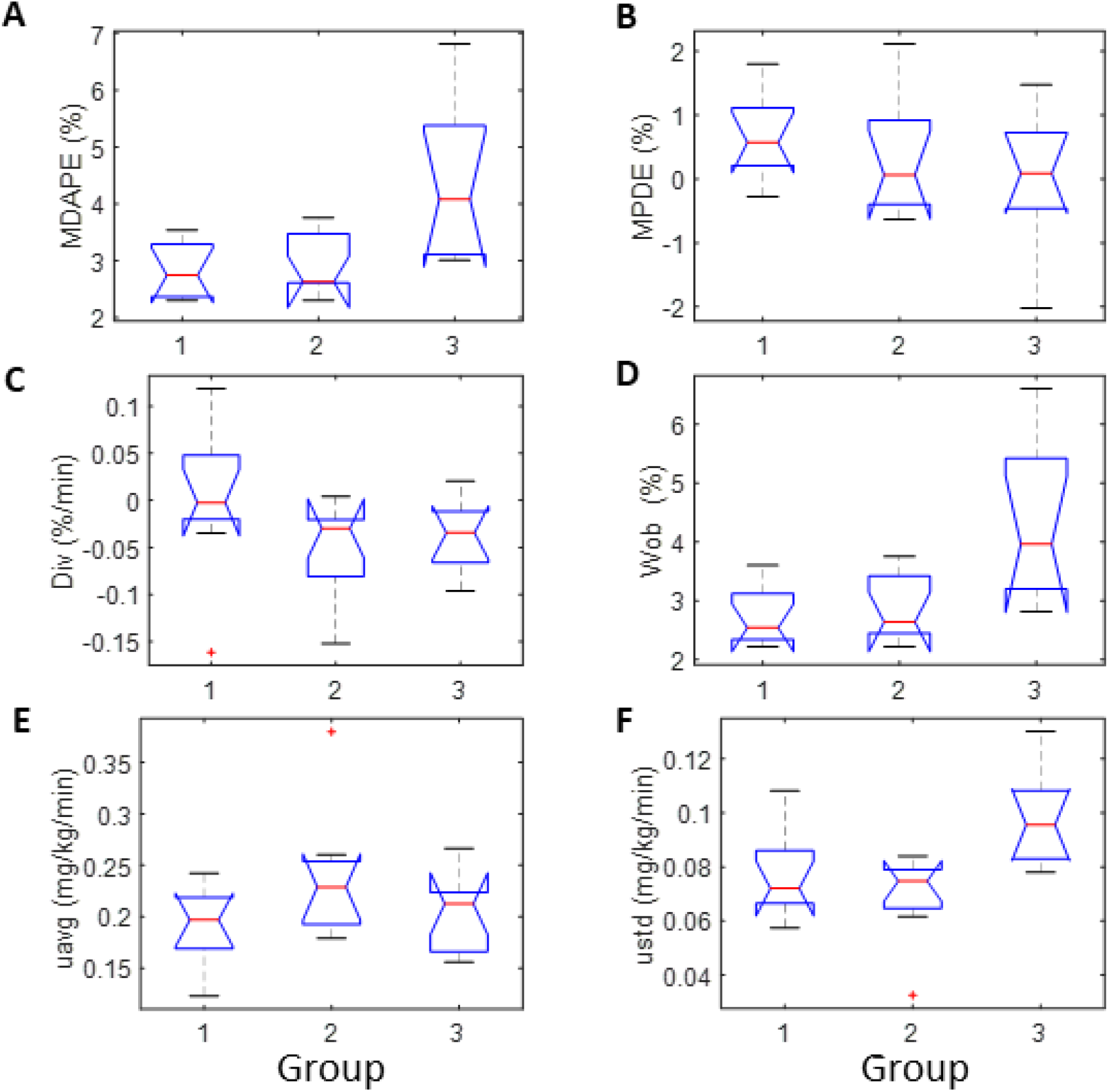
Box plots of performance measures across 2 NHPs and 2 target plans. Groups 1 through 3 respectively correspond to the three experimental designs: NHP-A with target plan 1, NHP-A with target plan 2, and NHP-B with target plan 1. The notches on the box plots correspond to the 95% CI on the median values for each metric.

## B Suggested future directions towards extending the current NHP CLAD study to future clinical studies

Although the CLAD system in this work is developed for the NHP model, the steps followed in this research can be adapted to develop a CLAD system for human applications using scalp EEG in place of LFPs.

The first step in this extension would be to choose a reliable EEG-based MOU whose temporal variation under propofol infusion shows similar trend as the 20-30 Hz LFP power in NHPs. Since the beta-frequency band activity in the EEG of adult volunteers under propofol anesthesia has similar trends to the NHP MOU [7], EEG-based MOUs that incorporate this information could be such candidate MOUs. Future NHP studies with simultaneous EEG, LFP and spiking activity recordings under propofol anesthesia can potentially provide a paradigm to connect the candidate EEG-based MOU to LFP and spiking activity. Neurophysiological modeling (such as [48, 49]) can provide a mechanistic basis for extrapolating neuroscientific interpretation of the EEG-based MOU in NHPs to EEG-based MOUs in humans. Furthermore, analysis of behavioral recordings simultaneously with EEG recordings in healthy volunteers (e.g. [7]) as well as patients (e.g. [50]) can further validate the candidate EEG-based MOUs as clinically-relevant biomarkers [51].

Once a reliable EEG-based MOU has been identified for humans, the next step would be to develop the corresponding PK-PD model. Unlike the limitation faced in the NHP studies, there exist several PK-PD models that have been used in human CLAD experiments or EEG monitoring studies [40, 28, 31, 11]. Many of these compartmental PK models are third or higher order ordinary differential equations. Our NHP CLAD studies show that a highly accurate anesthetic state control can be achieved with a second-order PK model. Given an EEG-based MOU for human studies, the parameters for the second-order human PK model with a sigmoidal PD model can be obtained by either allometric extrapolation [52] or by using the system identification approach we applied in our NHP studies (Sections 2.3, 4.4). Similarly, the control parameters can be computed using our tuning strategy in Section 2.4 (also Section 4.5).

Once a reliable PK-PD model for a reliable EEG-based MOU has been identified for humans, we can retrace the steps used in this work for feedback control design, and CLAD testing *in silico* and *in vivo* to develop a stable, accurate and robust CLAD system for human anesthesia care.

## Footnotes

1 Note: even in an ideal case when the true PK-PD parameters are available the model used in the control design can still be misspecified due to unaccounted nonlinearities (present in the true model) during the linearization step.

## References

[1] Thomas G Weiser, Scott E Regenbogen, Katherine D Thompson, Alex B Haynes, Stuart R Lipsitz, William R Berry, and Atul A Gawande. An estimation of the global volume of surgery: a modelling strategy based on available data. The Lancet, 372(9633):139–144, 2008.

[2] Burton S. Epstein. ASA Adopts Standards for the Practice of Anesthesiology. Archives of Surgery, 122(10):1215–1216, 10 1987.

[3] Denise Rohan, Donal J Buggy, Seamus Crowley, Ferraby KH Ling, Helen Gallagher, Ciaran Regan, and Denis C Moriarty. Increased incidence of postoperative cognitive dysfunction 24 hr after minor surgery in the elderly. Canadian Journal of Anesthesia, 52(2):137–142, 2005.

[4] Frederic A Gibbs, EL Gibbs, and WG Lennox. Effect on the electro-encephalogram of certain drugs which influence nervous activity. Archives of Internal Medicine, 60(1):154–166, 1937.

[5] Reginald G Bickford. Automatic electroencephalographic control of general anesthesia. Electroencephalography and Clinical Neurophysiology, 2(1-4):93–96, 1950.

[6] Donald K KIERSEY, REGINALD G Bickford, and Albert Faulconer Jr. Electro-encephalographic patterns produced by thiopental sodium during surgical operations: description and classification, 1951.

[7] Patrick L Purdon, Eric T Pierce, Eran A Mukamel, Michael J Prerau, John L Walsh, Kin Foon K Wong, Andres F Salazar-Gomez, Priscilla G Harrell, Aaron L Sampson, Aylin Cimenser, et al. Electroencephalogram signatures of loss and recovery of consciousness from propofol. Proceedings of the National Academy of Sciences, 110(12):E1142–E1151, 2013.

[8] Patrick L Purdon, Aaron Sampson, Kara J Pavone, and Emery N Brown. Clinical electroencephalography for anesthesiologists part i: background and basic signatures. Anesthesiology: The Journal of the American Society of Anesthesiologists, 123(4):937–960, 2015.

[9] H Schwilden, H Stoeckel, and J Schüttler. Closed-loop feedback control of propofol anaesthesia by quantitative eeg analysis in humans. BJA: British Journal of Anaesthesia, 62(3):290–296, 1989.

[10] Anthony R Absalom, Nicholas Sutcliffe, and Gavin N Kenny. Closed-loop control of anesthesia using bispectral indexper-formance assessment in patients undergoing major orthopedic surgery under combined general and regional anesthesia. Anesthesiology: The Journal of the American Society of Anesthesiologists, 96(1):67–73, 2002.

[11] Nicholas West, Guy A Dumont, Klaske van Heusden, Christian L Petersen, Sara Khosravi, Kristian Soltesz, Aryannah Umedaly, Eleanor Reimer, and J Mark Ansermino. Robust closed-loop control of induction and maintenance of propofol anesthesia in children. Pediatric Anesthesia, 23(8):712–719, 2013.

[12] Nicholas West, Klaske van Heusden, Matthias Görges, Sonia Brodie, Aryannah Rollinson, Christian L Petersen, Guy A Dumont, J Mark Ansermino, and Richard N Merchant. Design and evaluation of a closed-loop anesthesia system with robust control and safety system. Anesthesia & Analgesia, 127(4):883–894, 2018.

[13] Ngai Liu, Thierry Chazot, Antoine Genty, Alain Landais, Aymeric Restoux, Kathleen McGee, Pierre-Antoine Laloë, Bernard Trillat, Luc Barvais, and Marc Fischler. Titration of propofol for anesthetic induction and maintenance guided by the bispectral index: closed-loop versus manual control: a prospective, randomized, multicenter study. The Journal of the American Society of Anesthesiologists, 104(4):686–695, 2006.

[14] Ngai Liu, Thierry Chazot, Sophie Hamada, Alain Landais, Nathalie Boichut, Corinne Dussaussoy, Bernard Trillat, Laurent Beydon, Emmanuel Samain, Daniel I Sessler, et al. Closed-loop coadministration of propofol and remifentanil guided by bispectral index: a randomized multicenter study. Anesthesia & Analgesia, 112(3):546–557, 2011.

[15] Goverdhan D Puri, Preethy J Mathew, Indranil Biswas, Amitabh Dutta, Jayashree Sood, Satinder Gombar, Sanjeev Palta, Morup Tsering, PL Gautam, Aveek Jayant, et al. A multicenter evaluation of a closed-loop anesthesia delivery system: a randomized controlled trial. Anesthesia & Analgesia, 122(1):106–114, 2016.

[16] Maryam M Shanechi, Jessica J Chemali, Max Liberman, Ken Solt, and Emery N Brown. A brain-machine interface for control of medically-induced coma. PLoS computational biology, 9(10):e1003284, 2013.

[17] ShiNung Ching, Max Y Liberman, Jessica J Chemali, M Brandon Westover, Jonathan D Kenny, Ken Solt, Patrick L Purdon, and Emery N Brown. Real-time closed-loop control in a rodent model of medically induced coma using burst suppression. Anesthesiology: The Journal of the American Society of Anesthesiologists, 119(4):848–860, 2013.

[18] Bahram Parvinian, Christopher Scully, Hanniebey Wiyor, Allison Kumar, and Sandy Weininger. Regulatory considerations for physiological closed-loop controlled medical devices used for automated critical care: food and drug administration workshop discussion topics. Anesthesia and analgesia, 126(6):1916, 2018.

[19] Blackrock microsystems.

[20] T. Hagiwara, E. Furutani, and M. Araki. Two-degree-of-freedom design method of lqi servo systems: performance deterioration by the introduction of an observer and optimal observer design. In Proceedings of 1994 33rd IEEE Conference on Decision and Control, volume 4, pages 4204–4209 vol. 4, 1994.

[21] Tomomichi Hagiwara, Eiko Furutani, and Mituhiko Araki. Two-degree-of-freedom design method of linear-quadratic servo systems with an integral compensator: analysis of the performance deterioration by the introduction of an observer. International Journal of Control, 64(5):941–958, 1996.

[22] T. Hagiwara. Optimal observers for disturbance rejection in two-degree-of-freedom lqi servo systems. IEE Proceedings - Control Theory and Applications, 144:575–581(6), November 1997.

[23] Yumiko Ishizawa, Omar J Ahmed, Shaun R Patel, John T Gale, Demetrio Sierra-Mercado, Emery N Brown, and Emad N Eskandar. Dynamics of propofol-induced loss of consciousness across primate neocortex. Journal of Neuroscience, 36(29):7718–7726, 2016.

[24] Andre M Bastos, Jacob A Donoghue, Scott L Brincat, Meredith Mahnke, Jorge Yanar, Josefina Correa, Ayan S Waite, Mikael Lundqvist, Jefferson Roy, Emery N Brown, et al. Neural effects of propofol-induced unconsciousness and its reversal using thalamic stimulation. bioRxiv, 2020.

[25] Jesus Javier Ballesteros, Jessica Blair Briscoe, and Yumiko Ishizawa. Neural signatures of alpha2-adrenergic agonist-induced unconsciousness and awakening by antagonist. eLife, 9:e57670, aug 2020.

[26] Jesus J. Ballesteros, Pamela Huang, Shaun R. Patel, Emad N. Eskandar, and Yumiko Ishizawa. Dynamics of Ketamine-induced Loss and Return of Consciousness across Primate Neocortex. Anesthesiology, 132(4):750–762, 04 2020.

[27] Hartmut Derendorf and Bernd Meibohm. Modeling of pharmacokinetic/pharmacodynamic (pk/pd) relationships: concepts and perspectives. Pharmaceutical research, 16(2):176–185, 1999.

[28] Thomas W Schnider, Charles F Minto, Steven L Shafer, Pedro L Gambus, Corina Andresen, David B Goodale, and Elizabeth J Youngs. The influence of age on propofol pharmacodynamics. Anesthesiology: The Journal of the American Society of Anesthesiologists, 90(6):1502–1516, 1999.

[29] Sourish Chakravarty, Ksenia Nikolaeva, Devika Kishnan, Francisco J Flores, Patrick L Purdon, and Emery N Brown. Pharmacodynamic modeling of propofol-induced general anesthesia in young adults. In 2017 IEEE Healthcare Innovations and Point of Care Technologies (HI-POCT), pages 44–47. IEEE, 2017.

[30] Guy Chavent. Nonlinear least squares for inverse problems: theoretical foundations and step-by-step guide for applications. Springer Science & Business Media, 2010.

[31] Guy A Dumont, Arturo Martinez, and J Mark Ansermino. Robust control of depth of anesthesia. International Journal of Adaptive Control and Signal Processing, 23(5):435–454, 2009.

[32] Guy A Dumont and J Mark Ansermino. Closed-loop control of anesthesia: a primer for anesthesiologists. Anesthesia & Analgesia, 117(5):1130–1138, 2013.

[33] Sourish Chakravarty, Ayan S. Waite, John H. Abel, and Emery N. Brown. A simulation-based comparative analysis of pid and lqg control for closed-loop anesthesia delivery. IFAC-PapersOnLine, 53(2):15898–15903, 2020. 21th IFAC World Congress.

[34] David Luenberger. Observers for multivariable systems. IEEE Transactions on Automatic Control, 11(2):190–197, 1966.

[35] Michel M. R. F. Struys, Tom De Smet, Linda F. M. Versichelen, Stijn Van de Velde, Rudy Van den Broecke, and Eric P. Mortier. Comparison of Closed-loop Controlled Administration of Propofol Using Bispectral Index as the Controlled Variable versus “Standard Practice” Controlled Administration. Anesthesiology, 95(1):6–17, 07 2001.

[36] Emery N Brown, Kara J Pavone, and Marusa Naranjo. Multimodal general anesthesia: theory and practice. Anesthesia and analgesia, 127(5):1246, 2018.

[37] Ngai Liu, Morgan Le Guen, Fatima Benabbes-Lambert, Thierry Chazot, Bernard Trillat, Daniel I Sessler, and Marc Fischler. Feasibility of closed-loop titration of propofol and remifentanil guided by the spectral m-entropy monitor. The Journal of the American Society of Anesthesiologists, 116(2):286–295, 2012.

[38] Hemant Bokil, Peter Andrews, Jayant E Kulkarni, Samar Mehta, and Partha P Mitra. Chronux: a platform for analyzing neural signals. Journal of neuroscience methods, 192(1):146–151, 2010.

[39] Matthias Kreuzer. Eeg based monitoring of general anesthesia: Taking the next steps. Frontiers in Computational Neuroscience, 11:56, 2017.

[40] Thomas W Schnider, Charles F Minto, Pedro L Gambus, Corina Andresen, David B Goodale, Steven L Shafer, and Elizabeth J Youngs. The influence of method of administration and covariates on the pharmacokinetics of propofol in adult volunteers. Anesthesiology: The Journal of the American Society of Anesthesiologists, 88(5):1170–1182, 1998.

[41] Yuxiao Yang and Maryam M Shanechi. An adaptive and generalizable closed-loop system for control of medically induced coma and other states of anesthesia. Journal of neural engineering, 13(6):066019, 2016.

[42] H Schwilden. A general method for calculating the dosage scheme in linear pharmacokinetics. European Journal of Clinical Pharmacology, 20(5):379–386, 1981.

[43] T Miyabe-Nishiwaki, K Masui, A Kaneko, K Nishiwaki, T Nishio, and Hideko Kanazawa. Evaluation of the predictive performance of a pharmacokinetic model for propofol in japanese macaques (macaca fuscata fuscata). Journal of veterinary pharmacology and therapeutics, 36(2):169–173, 2013.

[44] Karl Johan Aström and Richard M Murray. Feedback systems: an introduction for scientists and engineers. Princeton university press, 2010.

[45] Robert H Shumway and David S Stoffer. Time series analysis and its applications: with R examples. Springer, 2017.

[46] John R Varvel, David L Donoho, and Steven L Shafer. Measuring the predictive performance of computer-controlled infusion pumps. Journal of pharmacokinetics and biopharmaceutics, 20(1):63–94, 1992.

[47] Michel M. R. F. Struys, Tom De Smet, Scott Greenwald, Anthony R. Absalom, Servaas Bingé, and Eric P. Mortier. Performance Evaluation of Two Published Closed-loop Control Systems Using Bispectral Index Monitoring: A Simulation Study. Anesthesiology, 100(3):640–647, 03 2004.

[48] Michelle M McCarthy, Emery N Brown, and Nancy Kopell. Potential network mechanisms mediating electroencephalographic beta rhythm changes during propofol-induced paradoxical excitation. Journal of Neuroscience, 28(50):13488–13504, 2008.

[49] Austin E Soplata, Michelle M McCarthy, Jason Sherfey, Shane Lee, Patrick L Purdon, Emery N Brown, and Nancy Kopell. Thalamocortical control of propofol phase-amplitude coupling. PLoS computational biology, 13(12):e1005879, 2017.

[50] John H Abel, Marcus A Badgeley, Benyamin Meschede-Krasa, Gabriel Schamberg, Indie C Garwood, Kimaya Lecamwasam, Sourish Chakravarty, David W Zhou, Matthew Keating, Patrick L Purdon, et al. Machine learning of eeg spectra classifies unconsciousness during gabaergic anesthesia. Plos one, 16(5):e0246165, 2021.

[51] FDA-NIH Biomarker Working Group et al. Best (biomarkers, endpoints, and other tools) resource [internet]. 2016.

[52] Donald E Mager, Sukyung Woo, and William J Jusko. Scaling pharmacodynamics from in vitro and preclinical animal studies to humans. Drug metabolism and pharmacokinetics, 24(1):16–24, 2009.

